# Nanoparticles with Curcumin and Piperine Modulate Steroid Biosynthesis in Prostate Cancer

**DOI:** 10.1101/2024.10.24.620019

**Authors:** Jibira Yakubu, Evangelos Natsaridis, Therina du Toit, Isabel Sousa Barata, Oya Tagit, Amit V. Pandey

## Abstract

Endogenous androgens are pivotal in the development and progression of prostate cancer (PC). We investigated nanoparticle formulations of curcumin and piperine in modulating steroidogenesis within PC cells. Using multiple PC cell lines (LNCaP, VCaP, DU145 and PC3) we studied the effects of curcumin, piperine, and their nanoparticle formulations—curcumin nanoparticles, piperine nanoparticles, and curcumin-piperine nanoparticles (CPN)—on cell viability, migration, and steroid biosynthesis. Curcumin and its nanoparticle formulations significantly reduced cell viability in PC cells, with curcumin-piperine nanoparticles showing the highest efficacy. These treatments also inhibited cell migration, with CPN exhibiting the most pronounced effect. In assays for steroid biosynthesis, curcumin and its nanoparticle formulations, as well as piperine and its nanoparticles, selectively inhibited 17α-hydroxylase and 17,20-lyase activities of cytochrome P450 17A1 (CYP17A1). Abiraterone, a CYP17A1 inhibitor, displayed a broader inhibition of steroid metabolism including cytochrome P450 21-hydroxylase activity, whereas curcumin and piperine provided a more targeted inhibition profile. Analysis of steroid metabolites by liquid chromatography-mass spectrometry revealed that CPN caused significant reduction of androstenedione and cortisol, suggesting potential synergistic effects. In conclusion, nanoformulations co-loaded with curcumin and piperine offer an effective approach to targeting steroidogenesis and could be promising candidates for therapies aimed at managing androgen-dependent PC.

## 1.0. Introduction

The prostatic microenvironment requires androgen receptor (AR) signaling to maintain normal homeostasis. Overactivation of AR signaling is a primary driver of prostate cancer (PC) development ^1,2^. Consequently, androgen deprivation is considered one of the most aggressive and effective initial treatments for malignant PC ^3,4^. Androgen deprivation therapy (ADT) is the cornerstone of treatment for advanced and metastatic PC (Figure 1). ADT functions primarily by suppressing or blocking testosterone, thus depriving the tumor of a crucial growth factor. Although ADT has improved disease-free survival when used as neoadjuvant or adjuvant therapy, it is associated with several adverse events, including gynecomastia, loss of libido, sexual dysfunction, hot flashes, sarcopenia, osteopenia, osteoporosis, metabolic syndrome, and cardiovascular-related issues, significantly impacting the quality of life of patients ^4,5^. Given that approximately one in nine men will be affected by PC in their lifetime, the number of individuals undergoing this treatment is substantial and growing ^6^.

**Figure 1.**
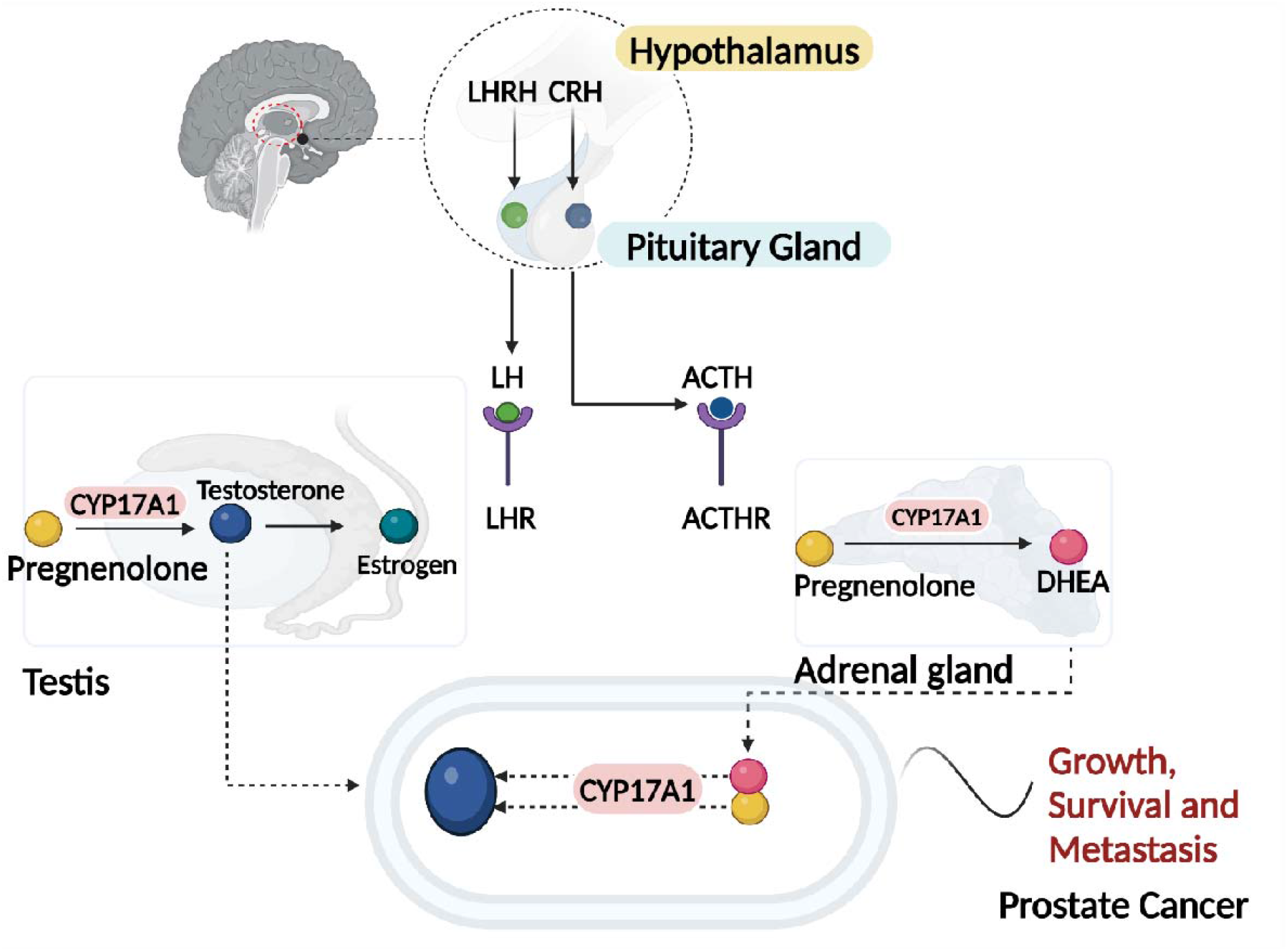
Androgen Production Pathways leading to the progression of Prostate Cancer, this diagram illustrates the hormonal regulation and production of androgens in the context of prostate cancer, highlighting the roles of the hypothalamus, testes, adrenal glands, and the prostate. Hypothalamus: The hypothalamus secretes luteinizing hormone-releasing hormone (LHRH) and corticotropin-releasing hormone (CRH). LHRH stimulates the anterior pituitary gland to release luteinizing hormone (LH), while CRH stimulates the release of adrenocorticotropic hormone (ACTH). In response to LHRH, the anterior pituitary releases LH, which acts on the Leydig cells in the testes to stimulate testosterone production. In response to CRH, the anterior pituitary releases ACTH, which stimulates the adrenal glands to produce androgens. LH binds to receptors (LHR) on Leydig cells, leading to the production and release of testosterone. Testosterone is the primary androgen and can be converted to the more potent dihydrotestosterone (DHT) by the enzyme 5α-reductase in target tissues, including the prostate. ACTH stimulates the adrenal cortex to produce androgens such as androstenedione and dehydroepiandrosterone (DHEA). These adrenal androgens can be converted to testosterone and other active androgens in peripheral tissues. Prostate cancer cells can utilize testosterone and DHT for growth and proliferation. Additionally, these cells can express enzymes that convert adrenal androgens to testosterone, maintaining androgen receptor signaling even when circulating testosterone levels are low. Created with Biorender.com.

To achieve androgen deprivation, medications such as luteinizing hormone-releasing hormone (LHRH) analogs, synthetic estrogens, gonadotropin-releasing hormone (GnRH) antagonists, AR blockers, and inhibitors of the enzyme cytochrome P450 17A1 (CYP17A1) involved in steroid synthesis ^7^, are used. Abiraterone is an example of an irreversible inhibitor of CYP17A1 and can reduce tumor burden, prolong life and relieve symptoms as it inhibits both the 17α-hydroxylase and 17,20-lyase activities of CYP17A1 and therefore inhibits multiple steps in androgen biosynthesis ^8,9^. However, resistance to the drug eventually allows disease progression ^10,11^.

Curcumin, a polyphenol found in the Curcuma longa plant, a spice extensively used as a culinary ingredient, has attracted attention due to its antioxidant, antimicrobial, and anti-inflammatory properties ^12^. Despite over 40 years of research into its medicinal properties, its mechanisms of action and exact molecular targets remain unclear. Previous data from our lab demonstrated that curcumin inhibits CYP17A1, and other cytochrome P450s involved in steroidogenesis. Our molecular docking studies also showed a high binding score for curcumin with CYP17A1 ^13^. Despite its well-documented effects, bioavailability of curcumin is compromised by poor absorption, rapid metabolism, and systemic elimination. Formulations such as polymeric nanoparticles, liposomes, phospholipid complexes, structural analogs, and derivatives have been investigated to overcome this hurdle ^12-15^. Poly (lactic-co-glycolic acid) (PLGA)-based nano-formulations, which are non-toxic, biodegradable, and non-immunogenic, hold value in medical applications ^16-18^. These clinical successes of PLGA formulations of have completed different clinical trial phases for cancer ^19,20^ and have inspired us to encapsulate curcumin in PLGA nanoparticles as a potential therapeutic option for androgen dependent PC. It has also been demonstrated that curcumin’s bioavailability can be improved by formulating it with PLGA polymer or PLGA combined with other polymers/copolymers ^14,17,21-24^.

Combinations of other phytochemicals with curcumin inhibits the progression of cancers, including PC ^12,25-27^. When a secondary active drug agent or drug candidate is co-administered with curcumin, an increase in the therapeutic benefit from curcumin has been reported in diverse cancer models. Notably, the second agent often enhances curcumin-dependent anti-cancer activity in a synergistic manner. A key candidate among these second agents is piperine, a dietary polyphenol isolated from black and long peppers. Piperine not only improves curcumin’s anti-cancer activity but also enhances its extremely poor bioavailability ^27-30^. Piperine alone displays anti-mutagenic and anti-tumor activities ^25,31,32^ inhibits the proliferation of colon cancer cell lines by inducing cell cycle arrest ^33^, and triggers apoptosis in osteosarcoma ^34^ and PC cells ^33^.

Effective therapies with limited side effect profiles based on scientifically justified rationales, such as phytochemicals, are urgently needed for many men diagnosed with PC, especially due to the disease’s progression and mortality rate. Here we show that a synergistic approach using curcumin and piperine as an inhibitor of the androgen signaling pathway may provide a more efficacious treatment option than curcumin alone in androgen sensitive PC. Therefore, we aimed to determine whether curcumin together with piperine can inhibit CYP17A1 activities in PC. We analyzed the effects of nanoencapsulated curcumin and piperine on the adrenal cell line NCI-H295R and multiple PC cell lines.

## Results and Discussion

### Colloidal and structural characterization of the nanoparticles

The hydrodynamic size and size distribution of empty and loaded PLGA nanoparticles were evaluated using DLS analysis. The average diameters of the prepared CN and CPN were found to be 177.2 ± 20.13 nm and 202.5 ± 36.33 nm, respectively, with corresponding polydispersity index (PDI) values of 0.091 ± 0.019 for CN and 0.122 ± 0.088 for CPN (Figure 2a). These measurements indicate a relatively uniform size distribution, particularly for CN. In comparison, the control nanoparticles and the piperine-only nanoparticles had diameters of 153.4 ± 16.05 nm and 169.3 ± 17.84 nm, respectively. The PDI values for these nanoparticles were 0.04 ± 0.017 for the control and 0.074 ± 0.019 for the piperine nanoparticles, suggesting a narrower size distribution than their curcumin-containing counterparts. The encapsulation efficiency (EE) of curcumin and piperine within the nanoparticles was also assessed. Curcumin demonstrated an encapsulation efficiency of 73.34 ± 4.32%, while piperine showed an encapsulation efficiency of 73.56 ± 6.78%. However, when co-encapsulated with piperine, the encapsulation efficiency of curcumin decreased by approximately 6%. This reduction in EE upon co-encapsulation aligns with previous reports of curcumin nanoparticles prepared using PLGA and other polymer matrices ^35-39^.

**Figure 2:**
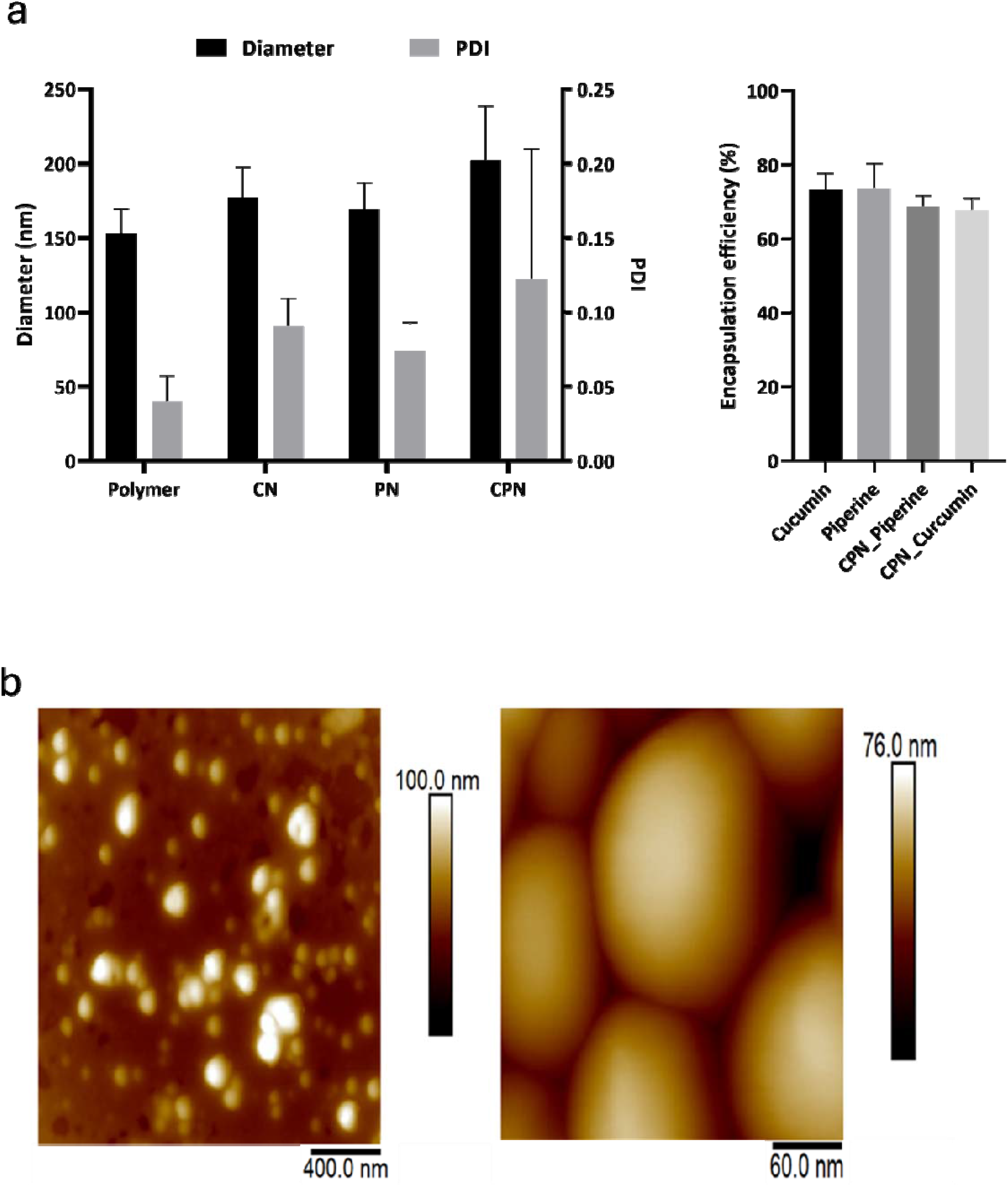
(a) Diameter and Polydispersity Index (PDI) of the prepared nanoparticles, including curcumin nanoparticles (CN), curcumin-piperine nanoparticles (CPN), and control nanoparticles. (b) Atomic Force Microscopy (AFM) height images of air-dried CPN with different scan sizes. Left panel; scan size: 2 μm x 2 μm. X-Y scalebar: 400 nm. Right panel; scan size: 300 nm x 300 nm. X-Y scalebar: 60 nm.

These findings indicate the efficiency of PLGA nanoparticles to encapsulate not only the individual drugs separately, but also both drugs within the same nanocarriers. While nanoencapsulation is potentially advantageous for enhanced therapeutic efficacy through improved cellular uptake, co-encapsulation of drug combinations can enhance the pharmacokinetic profile and augment the therapeutic efficacy of drugs as have been shown for curcumin and piperine in other studies ^27-29,40^.

The AFM analysis of CPN (Figure 2b) revealed the spherical morphology of the nanoparticles of approximately 150 nm in diameter, which is slightly smaller in comparison to DLS results. It is typical to observe such difference because DLS measures the hydrodynamic size, which takes the hydration layer into account as well. On the other hand, nanoparticles were air-dried for the AFM measurements, which further causes them to flatten under the AFM probe, leading to a reduced apparent height compared to their lateral size. This observation is consistent with the known behavior of polymer-based nanoparticles, which can deform under certain conditions, such as during the sample preparation and imaging processes ^36^.

Overall, the observed physicochemical properties of the nanoparticles, including size, PDI, morphology, and encapsulation efficiency, confirm their suitability for potential biomedical applications, particularly in drug delivery systems aimed at maximizing the bioavailability and effectiveness of curcumin and piperine.

### Fluorescence microscopy and the absorption of curcumin

Curcumin demonstrates intriguing photophysical and photochemical properties due to its structural composition featuring two symmetrically attached o-methoxy phenols via an α, β-unsaturated β-diketone linker. This structural motif also induces keto-enol tautomerism, facilitating the observation of curcumin’s intracellular uptake under a fluorescence microscope ^41^. As illustrated in Figure 3a, the fluorescence intensity in cells treated with CPN was significantly higher than in those treated with non-encapsulated curcumin at a concentration of 20 µM after 4 hours of incubation. This result indicates that the curcumin-piperine PLGA-loaded nanoparticles are more effective in enhancing the uptake of curcumin. Piperine has been reported in other studies to enhance the bioavailability of curcumin by inhibiting its metabolic breakdown and increasing its absorption in the gastrointestinal tract ^26-30,42,43^. Additionally, piperine augments the anticancer properties of curcumin by modulating several molecular targets and signaling pathways, thereby enhancing its therapeutic efficacy. Ability of piperine to inhibit enzymes involved in drug metabolism, such as cytochrome P450, and its potential to increase the permeability of the intestinal barrier, further contribute to the improved bioavailability and efficacy of curcumin ^40,44-46^. Therefore, the curcumin-piperine loaded PLGA nanoparticles showed higher cellular uptake of curcumin compared to non-encapsulated curcumin at the same dose, making them a promising candidate for cancer therapy. The Caco-2 cell line was employed in this study due to its relevance as a model for human intestinal absorption. Caco-2 cells, derived from human colorectal adenocarcinoma, spontaneously differentiate to form tight junctions, microvilli, and other characteristics of enterocytes upon reaching confluence, making them an ideal in vitro model for studying oral drug delivery and absorption ^47,48^. The absorption of curcumin, CN, and CPN into NCI-H295R cells was quantitatively analyzed following exposure to 20 μM concentrations of each compound for 4 hours. After extraction and analysis using UV/Visible spectrophotometry, the concentrations of curcumin, CN, and CPN found in the cells were determined to be 0.778 μM, 2.482 μM, and 4.245 μM, respectively. Curcumin exhibited the lowest intracellular concentration at 0.778 μM, while CN and CPN showed higher concentrations of 2.425 μM and 4.243 μM, respectively (Figure 3b). The enhanced uptake observed with CPN in Caco-2 and NCI-H295R cells suggests that these nanoparticles could potentially improve the oral bioavailability of curcumin, which is notoriously limited by poor solubility and rapid metabolism. Furthermore, the use of curcumin as a fluorescent probe in this study underscores its potential in tracking the cellular uptake and distribution of other non-fluorescent drug candidates. This method could be extended to investigate the intracellular behavior of various therapeutic compounds when co-formulated with curcumin in nanoparticle systems. The combination of curcumin with piperine not only enhances the bioavailability of curcumin but also exploits inherent anticancer properties of piperine. Piperine has been shown to inhibit enzymes that metabolize drugs, thereby increasing the concentration of curcumin in the bloodstream ^28^. Additionally, ability of piperine to modulate drug efflux transporters further contributes to the increased intracellular concentration of curcumin, enhancing its therapeutic efficacy ^44,45^. In summary, the use of curcumin-piperine loaded PLGA nanoparticles demonstrates a significant improvement in cellular uptake and bioavailability of curcumin, providing a promising strategy for PC therapy and a valuable tool for tracking other drug candidates in cellular studies. Supplementary Figure S1 presents the calibration curve for piperine, which serves as a reference for the analysis.

**Figure 3:**
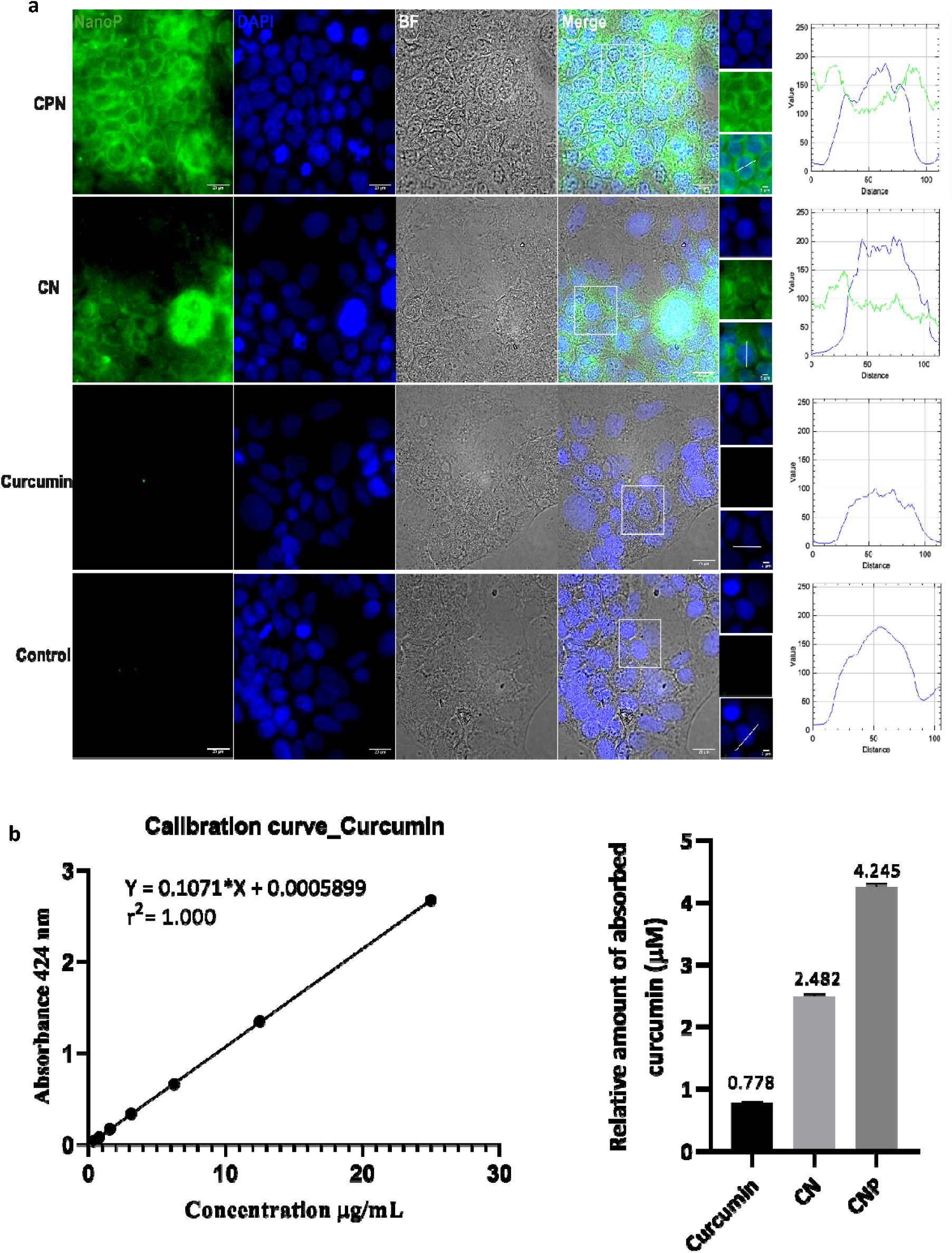
Absorption of nanoparticles. [a] Fluorescence microscopy images of Caco-2 cells after incubation with 20 µM curcumin at 37 °C for 4 hours. The images demonstrate the intracellular uptake and localization of curcumin loaded PLGA, highlighting its potential for cellular permeation and retention. Curcumin’s fluorescence properties allow for visualization within the cells, indicating its successful internalization. [b] Calibration curve of curcumin concentrations ranging from 0.391 to 25 µg/mL. The curve is used to determine the relative amount of curcumin absorbed by the cells, based on UV-vis absorbance intensity measurements. The strong linear correlation between curcumin concentration and absorbance intensity confirms the assay’s sensitivity and accuracy for quantifying intracellular curcumin levels. CN and CPN denotes curcumin nanoparticles, curcumin piperine nanoparticles respectively.

### Cytotoxicity of the drugs and nanoparticles on different PC cells

The cell viability assay conducted using the Alamar Blue assay revealed differential sensitivities of curcumin and its nanoparticle formulations across various PC cell lines, including androgen-sensitive (LNCaP and VCaP), androgen-insensitive (DU-145 and PC-3), and normal prostate epithelial cells (RWPE-1) (Figure 4). The Alamar Blue assay is a widely used, sensitive method for assessing cell viability and cytotoxicity. It is based on the reduction of resazurin, a non-toxic blue dye, to resorufin, a fluorescent pink compound, by mitochondrial enzymes in metabolically active cells ^49^. The intensity of fluorescence is directly proportional to the number of viable cells, providing a quantitative measure of cell viability. Curcumin and its nanoparticle formulations significantly reduced the viability of PC cell lines, with the curcumin-piperine nanoparticle formulation demonstrating the maximum efficacy, suggesting enhanced bioavailability and cellular uptake. Notably, curcumin-piperine nanoparticles selectively reduced viability in cancer cells while increasing viability in RWPE-1 cell, indicating potential for targeted therapy. Abiraterone and its nanoparticle formulation also decreased cell viability more effectively in androgen sensitive cell lines, with the nanoparticle form showing superior performance likely due to improved solubility and targeted delivery mechanisms. These findings align with existing literature that highlights the enhanced therapeutic efficacy of nanoparticle-based drug delivery systems ^16,20^. Nanoparticles have been shown to improve the solubility, stability, and bioavailability of anticancer agents, thereby increasing cytotoxicity in cancer cells while minimizing effects on normal cells ^50,51^. The selective cytotoxicity observed in this study aligns with reports that curcumin and its derivatives exhibit preferential toxicity towards PC cells through mechanisms such as apoptosis induction, oxidative stress, and inhibition of cell proliferation ^52-54^. Piperine has demonstrated notable anticancer properties against PC. It inhibits cancer cell proliferation by inducing cell cycle arrest at the G1 phase and promotes apoptosis through caspase activation and modulation of apoptotic proteins ^55,56^. Piperine also exerts anti-inflammatory and antioxidant effects by inhibiting the NF-κB pathway and reducing oxidative stress, contributing to its anticancer activity ^57^. A significant advantage of piperine is its ability to enhance the bioavailability of other compounds, such as curcumin, by inhibiting metabolic degradation. This results in increased systemic availability and effectiveness of curcumin ^28^. When used in combination, piperine and curcumin exhibit synergistic effects, leading to reduction of malignant effect of endocrine disrupting chemical. This combination also inhibits inflammatory signaling pathways critical for proliferative lesions ^31^. Furthermore, the enhanced effect of curcumin-piperine nanoparticles is consistent with findings that piperine enhances the bioavailability of curcumin by inhibiting its metabolic degradation ^28^. Previous work from our laboratory has demonstrated that doses below 10 µg/ml of curcumin have no toxic effect on HEK-293T (Supplementary Table S1) and adrenal NCI-H295R cells, highlighting its potential safety profile at lower concentrations ^13^.

**Figure 4.**
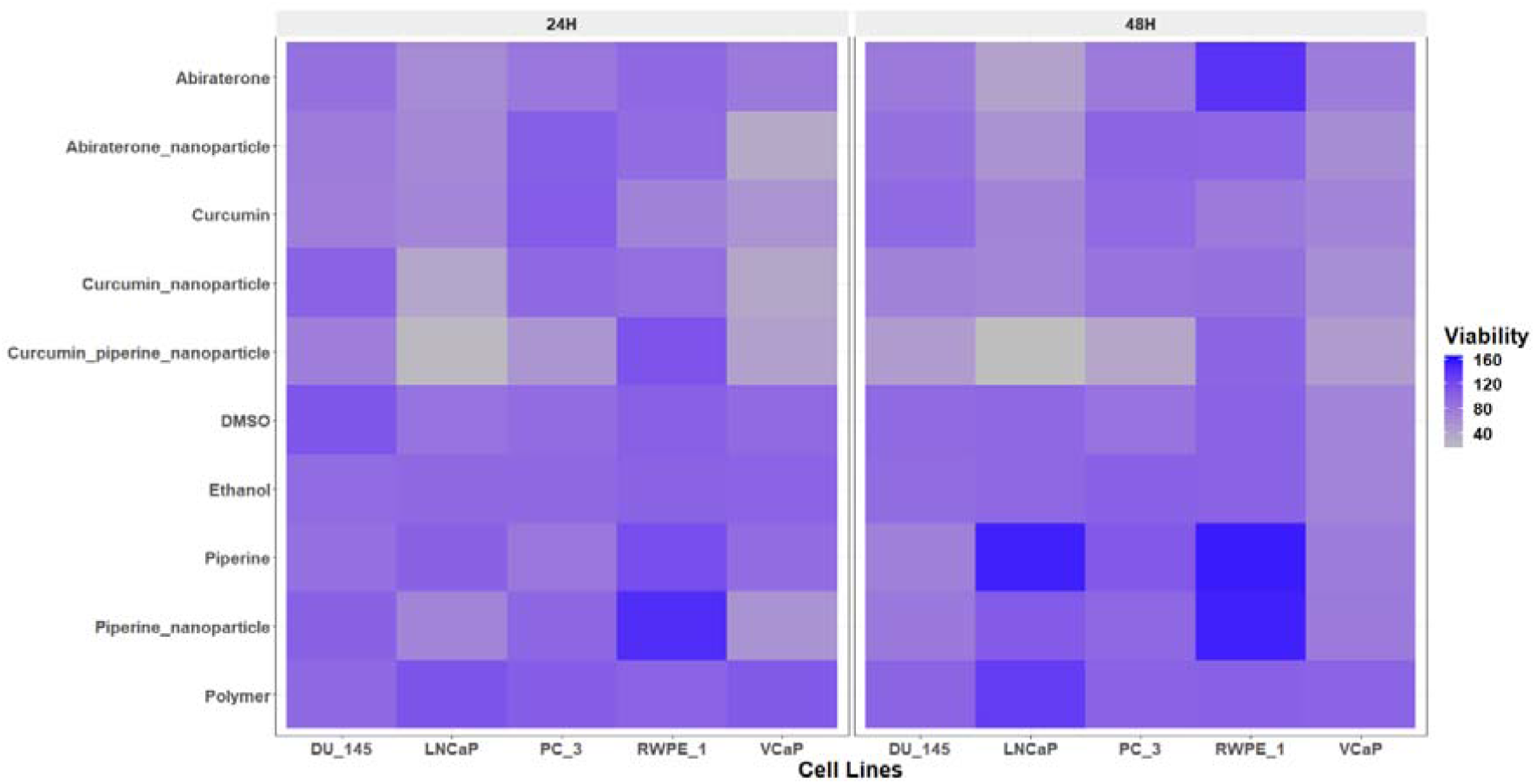
The heatmap illustrates the viability of PC cell lines (LNCaP, VCaP, DU--145, PC-3) and normal prostate epithelial cells (RWPE-1) following 24-hour and 48-hour treatments with different compounds: ethanol, polymer, DMSO, abiraterone, abiraterone nanoparticles, curcumin, curcumin nanoparticles, curcumin piperine nanoparticles, and piperine nanoparticles. Cell viability was assessed using the Alamar Blue assay and is represented as a percentage of the untreated control cells. The data points for each cell line and time point are averaged from triplicate experiments. The results highlight the differential sensitivities of the cell lines to the treatments, with curcumin piperine nanoparticles showing significant reduction in viability in PC cell lines, especially in the 48-hour treatments, while maintaining higher viability in normal RWPE-1 cell. The nanoparticle formulations of both curcumin and abiraterone exhibited enhanced efficacy compared to their free drug counterparts. Low viability (indicating higher cytotoxicity) is depicted in grey and high viability (indicating lower cytotoxicity) is depicted in blue.

### Wound healing effects of the drugs and the nanoparticles

The scratch assay results, expressed as the percentage of control, are shown in Figure 5. The polymer, DMSO, and ethanol controls showed no significant differences in cell migration. Abiraterone treatment resulted in a modest decrease in cell migration compared to the polymer control, although this difference was not statistically significant. Significant reduction in cell migration were observed with treatments involving curcumin, piperine, and their nanoparticle formulations (Supplementary Figure S3). Specifically, curcumin alone decreased cell migration significantly (mean difference = −183.3, p < 0.0001), as did piperine (mean difference = −125.7, p = 0.0015). Notably, CN and CPN formulations showed the best reductions in cell migration, with mean differences of −200.8 (p < 0.0001) and −203.7 (p < 0.0001), respectively, compared to the polymer control. When comparing abiraterone to curcumin and its formulations, significant reductions in cell migration were observed for curcumin (mean difference = −115.0, p = 0.0030), CN (mean difference = −132.5, p = 0.0010), and CPN (mean difference = −135.4, p = 0.0008). Finally, comparisons between piperine and its nanoparticle formulations showed that the CPN significantly reduced cell migration compared to piperine alone (mean difference = −77.95, p = 0.0422). Curcumin significantly inhibited cell migration, showing a marked reduction compared to the control, with an adjusted p-value < 0.0001. This result is consistent with previous studies indicating curcumin’s ability to inhibit PC cell proliferation and migration by down-regulating androgen receptors and epidermal growth factor receptors, inducing cell cycle arrest, and inhibiting NF-κB signaling. Piperine alone also significantly reduced cell migration compared to the control, with an adjusted p-value of 0.0015. This supports existing literature that piperine enhances curcumin’s bioavailability and synergistically increases its anti-cancer efficacy. CN and CPN exhibited the highest reduction in cell migration, which was statistically significant (adjusted p-value < 0.0001). This aligns with previous observations from our cytotoxicity study that CN and CPN has superior cytotoxic effects due to enhanced absorption and targeted delivery. PLGA nanoparticle delivery systems improve curcumin’s solubility, stability, and cellular uptake, leading to increased therapeutic efficacy. Our results demonstrate that curcumin and piperine, particularly in nanoparticle formulations, significantly inhibit PC cell migration. This suggests a promising therapeutic strategy for PC treatment, potentially preventing metastasis and improving patient outcomes. The micrographs from the scratch assay is represented as supplementary Figure S3.

**Figure 5:**
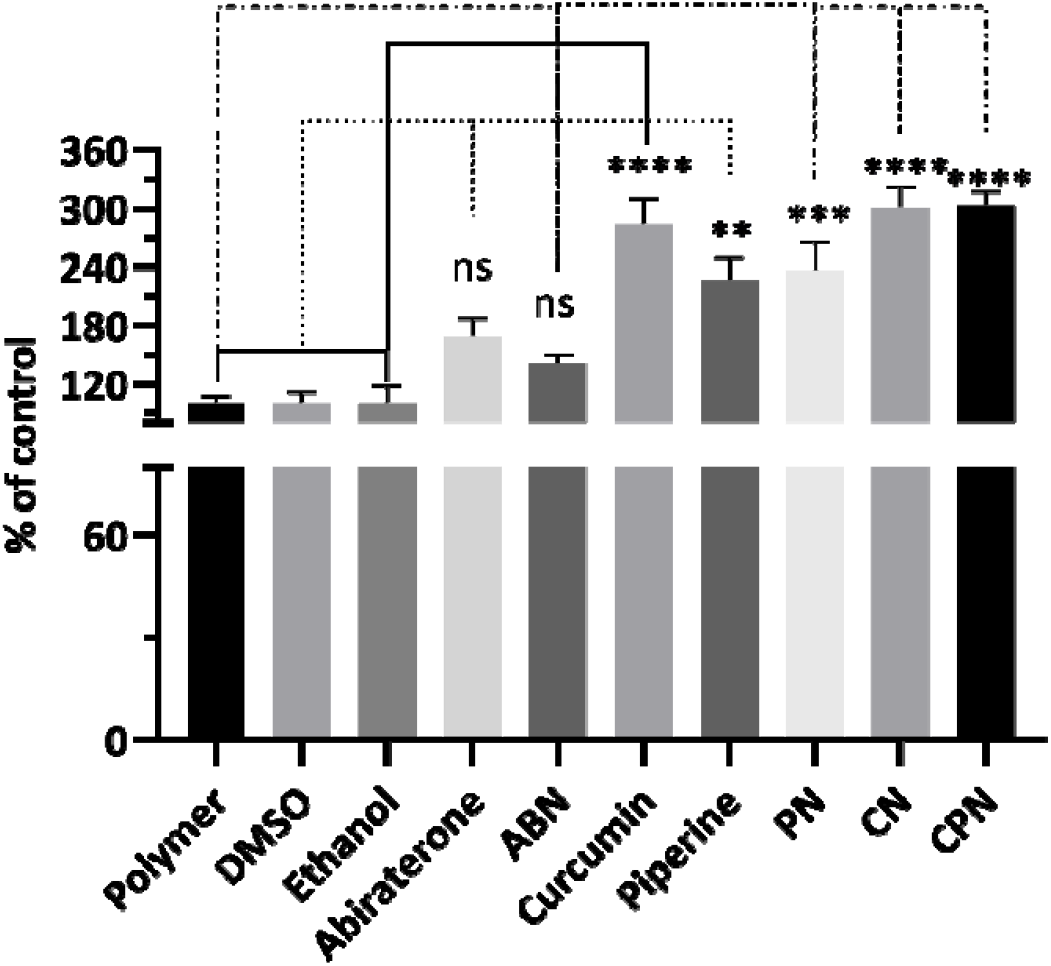
Analysis of cell migration by treatment with different compounds: Polymer, DMSO, Ethanol, Abiraterone, ABN (Abiraterone Nanoparticles), Curcumin, Piperine, PN (Piperine Nanoparticles), CN (Curcumin Nanoparticles), and CPN (Curcumin Piperine Nanoparticles). Cells were treated for 24 hours, and the migration was quantified as a percentage of the control. Data are presented as mean ± standard deviation (SD). Statistical analysis was performed using Tukey’s multiple comparisons test. Significant differences are indicated as follows: p < 0.01 (**), p < 0.001 (***), and p < 0.0001 (****), and ns, not significant

### Effects of drugs on steroid biosynthesis

The NCI-H295R cell line is instrumental in this assay due to its ability to express all steroidogenic enzymes found in the human adrenal cortex, making it an excellent model for studying steroid biosynthesis and its regulation ^8,58-60^. Using radioactive substrates, such as progesterone (^14^C) and 17α-hydoxypregnenolone (^3^H), allows for precise measurement of enzyme activity, providing robust and quantifiable data. Our study investigated the effects of curcumin, piperine, and their nanoparticle formulations on CYP17A1 hydroxylase and lyase activities in NCI-H295R cells, using DMSO, ethanol, and polymer as controls, with abiraterone as a positive control. Hydroxylase activity (Figure 6 a & b) remained close to 100% for DMSO, ethanol, and polymer controls, while abiraterone and its nanoparticles significantly inhibited this activity to 7.9% and 4.6%, respectively. Curcumin and its nanoparticles reduced hydroxylase activity to 76% and 65%, respectively. Piperine and its nanoparticles decreased hydroxylase activity to 87% and 88%, respectively. The combination of curcumin and piperine resulted in a significant reduction to 7.6%. For lyase activity (Figure 6c), controls maintained over 94%, with abiraterone and its nanoparticles inhibiting it to 13% and 14%. Curcumin and its nanoparticles inhibited lyase activity to 36% and 11%, respectively. Piperine and its nanoparticles showed inhibition of 81% and 71%, respectively, while their combination led to a notable reduction to 6%. The IC_50_ for curcumin nanoparticles on 17α-hydroxylase inhibition was 9.98 µM (Figure 6d). Abiraterone, a potent CYP17A1 inhibitor, is commonly used in the treatment of PC due to its ability to reduce androgen production. However, its use is associated with several limitations. Abiraterone can cause significant side effects, including hypertension, hypokalaemia, and liver toxicity ^61-64^. Additionally, its inhibition of CYP21A2 can lead to reduced cortisol production, necessitating concomitant corticosteroid therapy to manage side effects ^64^. Furthermore, resistance to abiraterone can develop, limiting its long-term efficacy in treating prostate cancer ^65,66^. These findings have significant scientific relevance as they underscore the potential of curcumin and piperine, especially in nanoparticle formulations, as effective inhibitors of CYP17A1 activities. The stronger inhibition of lyase activity compared to hydroxylase activity suggests a targeted approach in disrupting androgen biosynthesis, which is crucial in PC progression. The nanoparticle formulations of curcumin and piperine offer several advantages over abiraterone. Nanoparticles can enhance the bioavailability and stability of curcumin and piperine, ensuring more efficient delivery to target sites. This targeted delivery can potentially reduce side effects and improve therapeutic outcomes.

**Figure 6.**
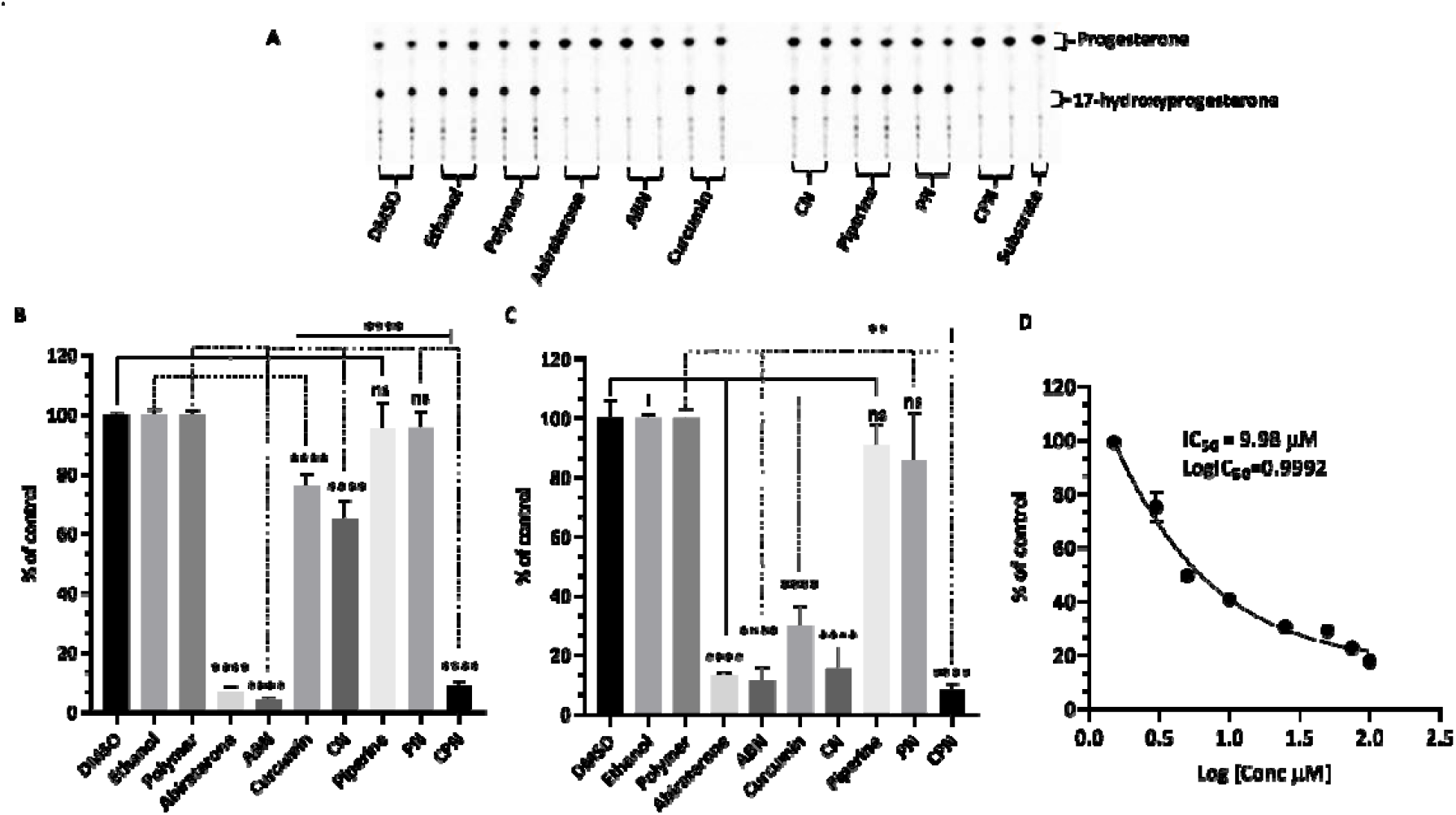
Assays of CYP17A1 activity. [a] Representative TLCs illustrating the effect of various treatments on CYP17A1 hydroxylase activity in NCI-H295R cells. Human adrenal NCI-H295R cells were treated with polymer, DMSO, ethanol, abiraterone, abiraterone nanoparticles (ABN), curcumin, piperine (Pip), piperine nanoparticles (PN), curcumin nanoparticles (CN), and curcumin piperine nanoparticles (CPN). Subsequently, cells were incubated with C-14 labelled progesterone, a substrate for CYP17A1-hydroxylase. Steroids were then extracted using organic solvents, dried under nitrogen, and separated by TLC. The radioactivity of the steroids was visualized using autoradiography on a phosphorimager and quantitated by densitometric analysis. [b] Curcuminoids inhibited the formation of 17α-hydroxyprogesterone in NCI-H295R cells, indicating that Cur & Pip caused significant inhibition of the 17α-hydroxylase. [c] Differential inhibition of various treatments on CYP17A1 17,20 lyase activity. Cells were treated with tritium-labelled 17 α-hydroxypregnenolone, and the tritiated by-product (acetate) was measured using a scintillation counter after DHEA was precipitated in a charcoal-dextran solution. Ethanol, phosphate buffer-polymer, and DMSO were used as controls at 0.1% of the total reaction volume. (d) Shows the CYP17A1 hydroxylase IC_50_ curve for the curcumin nanoparticles in NCI-H295R Cells. The statistical analysis was performed using Tukey’s multiple comparisons test. Significant differences are indicated as follows: p < 0.01 (**), p < 0.001 (***), and p < 0.0001 (****), ns, not significant.

### Effects of Compounds on CYP21A2 Activity

In human adrenal glands, CYP17A1 primarily catalyzes the 17α-hydroxylation of pregnenolone and progesterone, followed by a 17,20-lyase reaction of 17α-hydroxypregnenolone to produce dehydroepiandrosterone (DHEA) ^67^. This enzyme does not use 17α-hydroxyprogesterone as a substrate in humans, which is a key distinction from CYP21A2. CYP21A2 specifically catalyzes the hydroxylation of 17α-hydroxyprogesterone to produce 11-deoxycortisol, a crucial intermediate in the biosynthesis of glucocorticoids and cortisol. This substrate specificity allows for the precise assay of CYP21A2 activity by monitoring the conversion of 17α-hydroxyprogesterone to 11-deoxycortisol. The use of 17α-hydroxyprogesterone as a substrate is essential in distinguishing the enzymatic activities of CYP17A1 and CYP21A2. The reaction was monitored by measuring the production of 11-deoxycortisol, which serves as a direct indicator of CYP21A2 activity. This method leverages the specificity of 17α-hydroxyprogesterone for CYP21A2, ensuring accurate assessment of the enzyme’s function without interference from CYP17A1 activity. Cells were seeded overnight and then incubated with compounds for 4h. The effects of these compounds on CYP21A2 activity were assessed by measuring the conversion of [3H] 17α-hydroxyrogesterone to 11-deoxycortisol in NCI-H295R cells (Figure 7a&b). Controls with DMSO, ethanol and polymer-maintained enzyme activities close to 100%, confirming their suitability as solvents in these assays. Abiraterone, a known inhibitor, significantly reduced CYP21A2 activity to 8% of control, mirroring its inhibition of CYP17A1. This reduction indicates non-specific inhibition across steroidogenic enzymes, which is a critical consideration in therapeutic contexts where selective enzyme targeting is preferred. Curcumin reduced CYP21A2 activity to 83%, while curcumin nanoparticles exhibited a lesser inhibitory effect, maintaining 97% activity. Piperine had 86% of control activity in CYP21A2 assays, but its nanoparticle form interestingly increased enzyme activity to 111%. The combination of curcumin and piperine resulted in a slight increase in CYP21A2 activity to 111%, suggesting a potential interactive effect that mitigates inhibition. These findings highlight the differential impacts of curcumin and piperine on CYP21A2 activity compared to their stronger inhibition of CYP17A1. This selective inhibition profile is significant for therapeutic applications, particularly in PC treatment, where targeted disruption of androgen biosynthesis via CYP17A1 inhibition is desired without adversely affecting other steroidogenic pathways such as those mediated by CYP21A2. The use of nanoparticle formulations could enhance the specificity and efficacy of these compounds, offering a promising avenue for developing new treatments that mitigate the limitations of current therapies like abiraterone.

**Figure 7.**
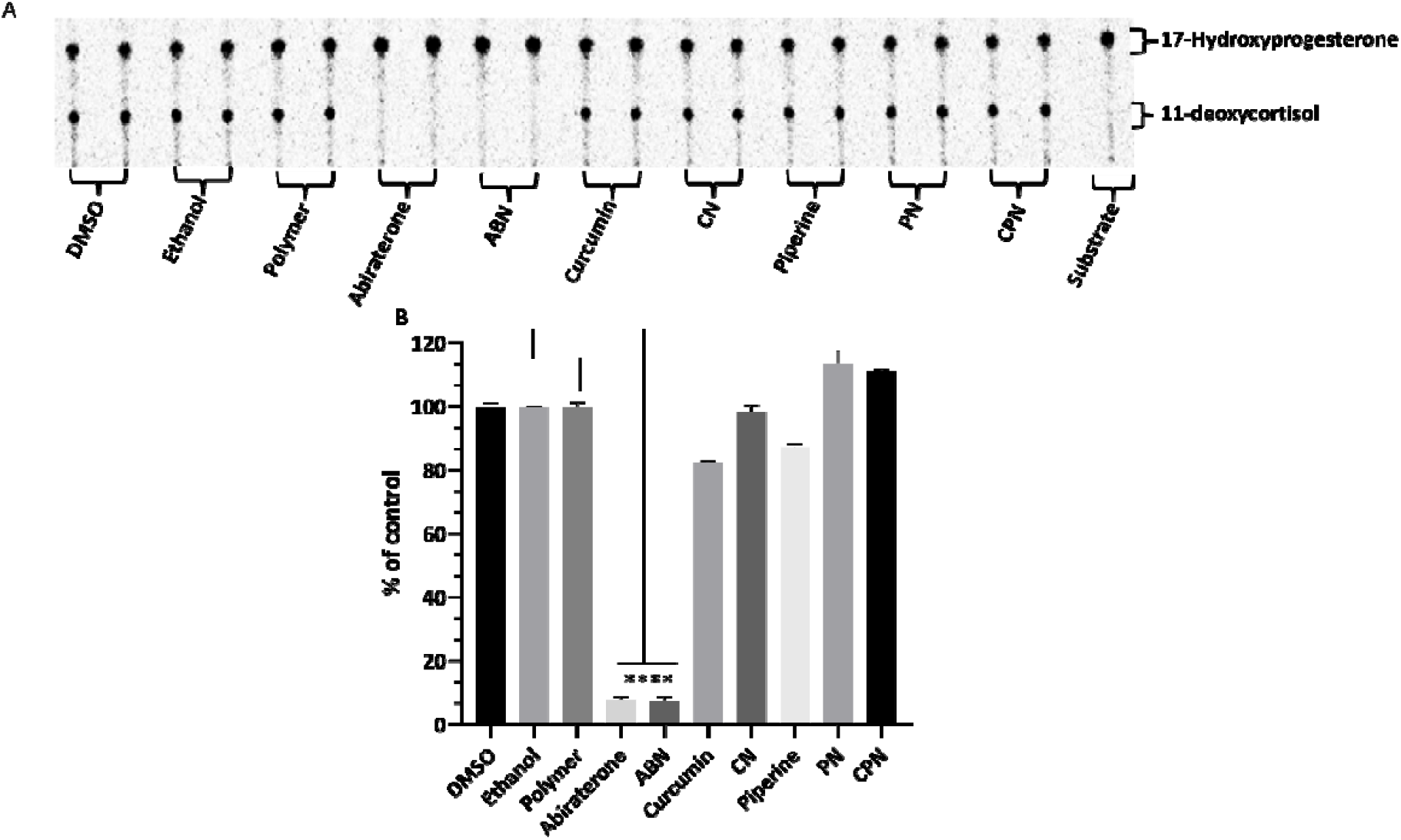
Assessment of CYP21A2 activity in human adrenal NCI-H295R cells. NCI-H295R cells were incubated for 4 hours with various treatments: polymer, DMSO, ethanol, abiraterone, ABN (abiraterone nanoparticles), curcumin, piperine, PN (piperine nanoparticles), CN (curcumin nanoparticles), and CPN (curcumin piperine nanoparticles). The activity of CYP21A2 was determined using [3H]-17α-hydroxyprogesterone as the substrate. Following incubation, steroids were extracted, dried, and separated by TLC. The conversion of 17α-hydroxyprogesterone to 11-deoxycortisol was visualized through autoradiographic analysis on TLC plates and quantified using densitometry. One-way ANOVA was employed to statistically compare each treatment group to the control, with significance indicated as **** p < 0.0001.

### LC-MS measurement of secreted steroid metabolites in the human adrenal H295R cells

We evaluated the effects of curcumin, curcumin nanoparticles (CN), curcumin-piperine nanoparticles (CPN), and the respective controls (DMSO, ethanol, polymer, and abiraterone) on secreted metabolites in NCI-H295R cells following 24 hours of treatment. The drugs were selected based on previous data indicating that piperine and its nanoparticles did not exhibit significant inhibitory effects on 17α-hydroxylase and lyase activity. CYP21A2, an enzyme crucial for producing glucocorticoids and mineralocorticoids, plays a pivotal role in cortisol and aldosterone synthesis, which is essential for maintaining metabolic homeostasis and regulating the hypothalamic-pituitary-adrenal (HPA) axis. Our data, presented in Table 1, reveal that abiraterone significantly decreased the production of several steroid metabolites, including androstenedione, cortisol, and testosterone, suggesting that its effects are not entirely selective for CYP17A1, as it also markedly inhibits CYP21A2 activity. The observed cross-inhibition of CYP21A2 by abiraterone could be attributed to the structural similarities between CYP17A1 and CYP21A2, even though these enzymes have relatively low sequence identity^64^. Structural homology between the active sites of these enzymes may allow abiraterone to bind to CYP21A2, thereby inhibiting its activity. Evidence from studies on other cytochrome P450 enzymes supports the notion that minor differences in protein sequences can still result in significant overlap in substrate and inhibitor binding capabilities due to conserved structural motifs within the enzyme family. The inhibition of CYP21A2, evidenced by the drastic reduction in 11-deoxycortisol and cortisol levels, highlights potential off-target effects that complicate its use in clinical settings. Curcumin and its nanoparticle formulation demonstrated notable effects on steroid metabolism. Curcumin significantly reduced levels of androstenedione and cortisol while preserving or slightly enhancing levels of other metabolites like progesterone and pregnenolone. This selective inhibition could reflect a targeted mechanism of action that spares some pathways while disrupting others. Notably, CPN exhibited a pronounced reduction in 17α-hydroxyprogesterone, aligning with previous findings on curcumin’s inhibitory effects on CYP17A1. The combination of curcumin and piperine resulted in a more substantial impact on steroid metabolism compared to curcumin nanoparticles alone. This observation suggests a potential synergistic interaction between curcumin and piperine, which may enhance their collective efficacy. LC-MS analysis data (Table 1) reveal that curcumin significantly decreased androstenedione and cortisol levels, and this effect was further amplified by the addition of piperine, as seen in the CPN treatment. This suggests that the combination of curcumin and piperine might work synergistically to more effectively target steroidogenic pathways. Enhanced efficacy of the CPN treatment compared to curcumin nanoparticles could be attributed to the complementary actions of the two compounds, which might synergistically inhibit multiple steps in steroid biosynthesis. The pronounced effects of the curcumin-piperine combination highlight its potential as a therapeutic strategy for managing adrenal-related conditions, demonstrating that combinations of bioactive compounds can offer more substantial effects than single-agent treatments. Additionally, these findings emphasize the importance of evaluating compound interactions and off-target effects in drug development to optimize therapeutic outcomes and minimize adverse effects.

**Table 1:**
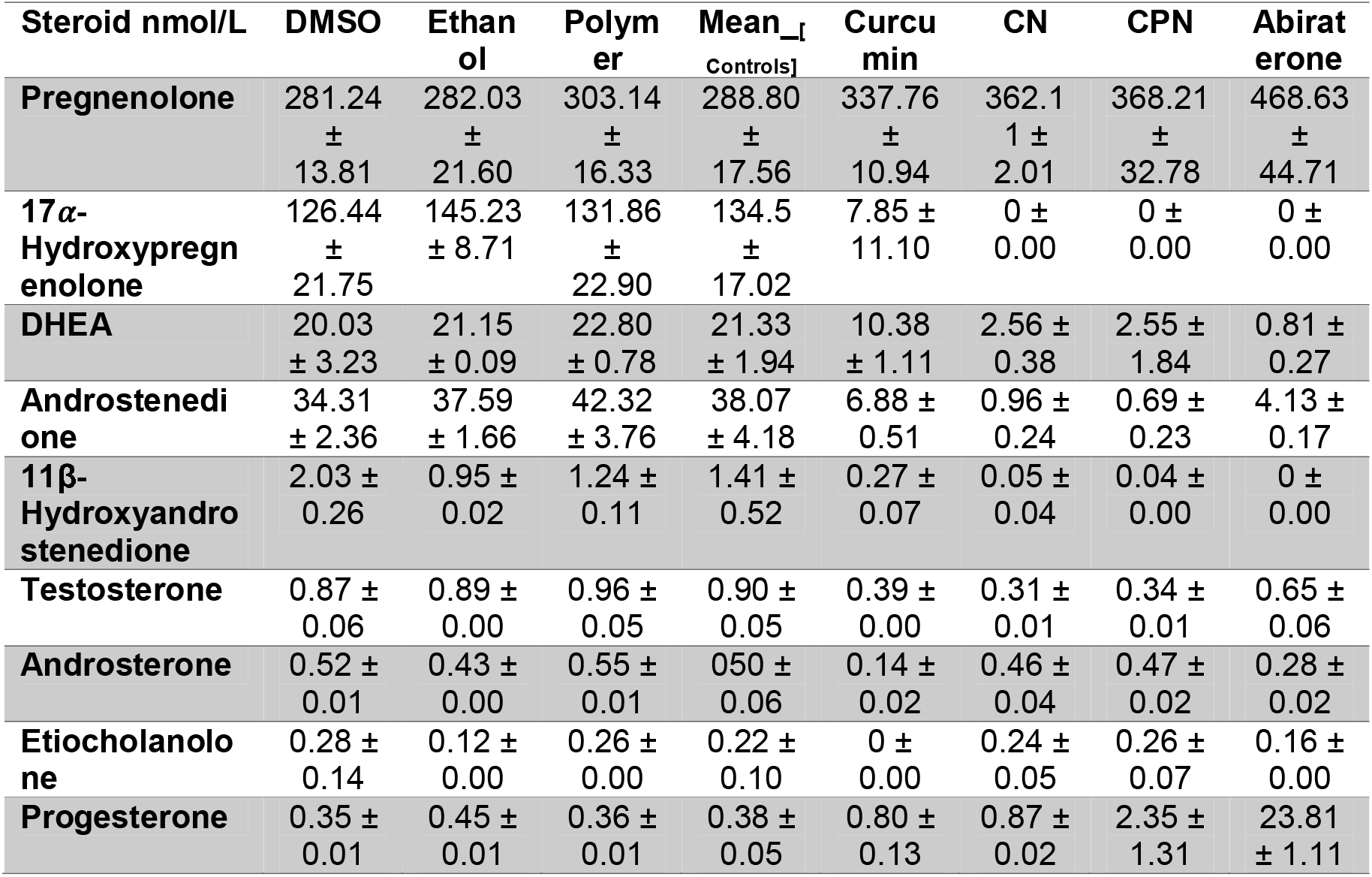
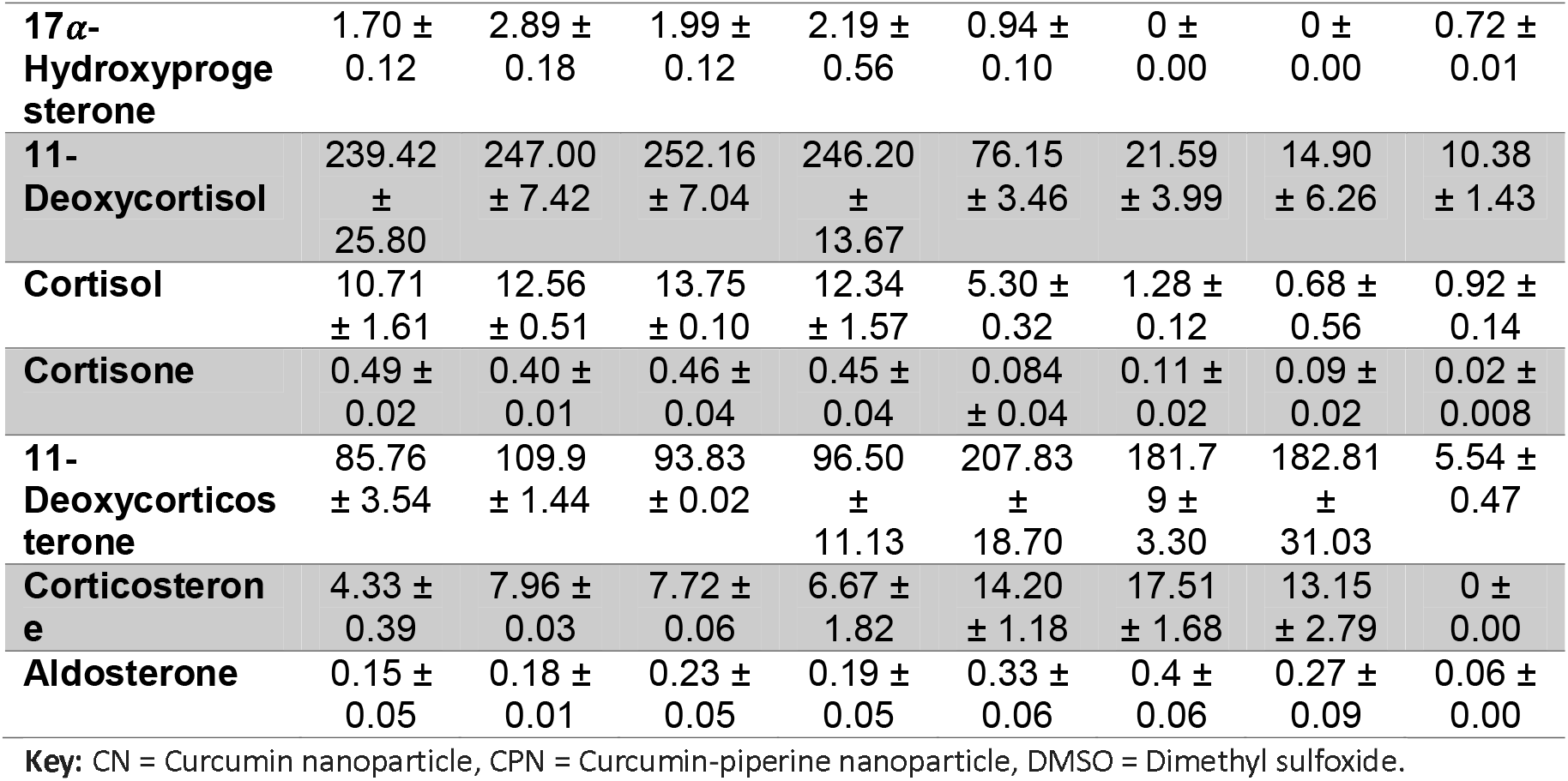
Steroid profiling in human adrenal NCI-H295R cells using LC-MS. Secreted steroid metabolites (measured in nmol/L) following treatment with pregnenolone (1 μM) and various compounds (10 μM) are shown (values in italics shows levels detected below the limit of accurate quantification: DHEA-S, DHT and 21-deoxycortisol were not detected).

### Effects of compounds on steroidogenic genes

Our study investigated the impact of various inhibitors on the gene expression of key enzymes involved in steroid biosynthesis in human adrenal NCI-H295R cells. We specifically assessed the effects of curcumin, piperine, and their nanoparticle formulations, alongside abiraterone, on the expression levels of CYP17A1, POR (P450 Oxidoreductase), and CYB5 (Cytochrome b5). These genes are critical components of the steroidogenic pathway, influencing the synthesis of androgens and other steroid hormones, which have profound implications in PC progression. CYP17A1, POR, and CYB5 play crucial roles in steroidogenesis, influencing androgen production and, consequently, PC progression ^62,68^. CYP17A1 is pivotal due to its dual enzymatic activities: 17α-hydroxylase and 17,20-lyase. The 17α-hydroxylase activity catalyzes the hydroxylation of pregnenolone and progesterone to form 17α-hydroxy pregnenolone and 17α-hydroxyprogesterone, respectively. Subsequently, the 17,20-lyase activity converts these intermediates into androstenedione and DHEA, essential precursors for the biosynthesis of androgens like testosterone and dihydrotestosterone. Elevated CYP17A1 activity leads to increased androgen levels, which can enhance tumor growth and resistance to therapies in prostate cancer, making its inhibition a strategic approach to lower androgen levels and impede tumor progression. POR is a critical electron donor for CYP450 enzymes, including CYP17A1, facilitating the transfer of electrons from NADPH to these enzymes, which is crucial for their catalytic activities. Activity of POR is integral to maintaining steroidogenic efficiency, and alterations in its expression can impact overall steroid biosynthesis and androgen production. Manipulating POR expression may affect the efficacy of CYP17A1 inhibitors and thus influence androgen levels ^67,69,70^. Similarly, CYB5 functions as an electron transfer protein that modulates steroidogenesis by enhancing the catalytic efficiency of CYP450 enzymes, including CYP17A1. CYB5 interacts with CYP17A1 to facilitate the conversion of 17α-hydroxy pregnenolone to DHEA, underscoring its role in regulating androgen levels. Variations in CYB5 expression can affect steroidogenic output and potentially influence PC progression, with reduced CYB5 levels potentially leading to decreased androgen production and impacting tumor growth ^71-73^.

### CYP17A1 Expression

CYP17A1, a crucial enzyme in androgen biosynthesis, was evaluated across different treatments. Abiraterone, a well-known CYP17A1 inhibitor, showed a slight increase in CYP17A1 expression, with values ranging from 1.13 to 1.19-fold of the control (Figure 8). In contrast, nanoparticle formulations of curcumin and piperine demonstrated varying degrees of inhibition on CYP17A1 expression. Curcumin nanoparticle formulation reduced CYP17A1 expression to 0.74-fold of abiraterone and 0.74-fold of control in different replicates Similarly, the nanoparticle formulation of piperine also reduced CYP17A1 expression, to 0.85-fold of abiraterone, though this was less pronounced than curcumin. Curcumin alone significantly decreased CYP17A1 expression to 0.74-fold of abiraterone and 0.70-fold of control, while piperine alone had a more enhanced effect, increasing CYP17A1 expression to 1.24-fold of abiraterone and 1.22-fold of control. These data shows that both curcumin and piperine, particularly in their nanoparticle formulations, exert regulatory effects on CYP17A1, supporting their role in modulating steroid biosynthesis.

**Figure 8:**
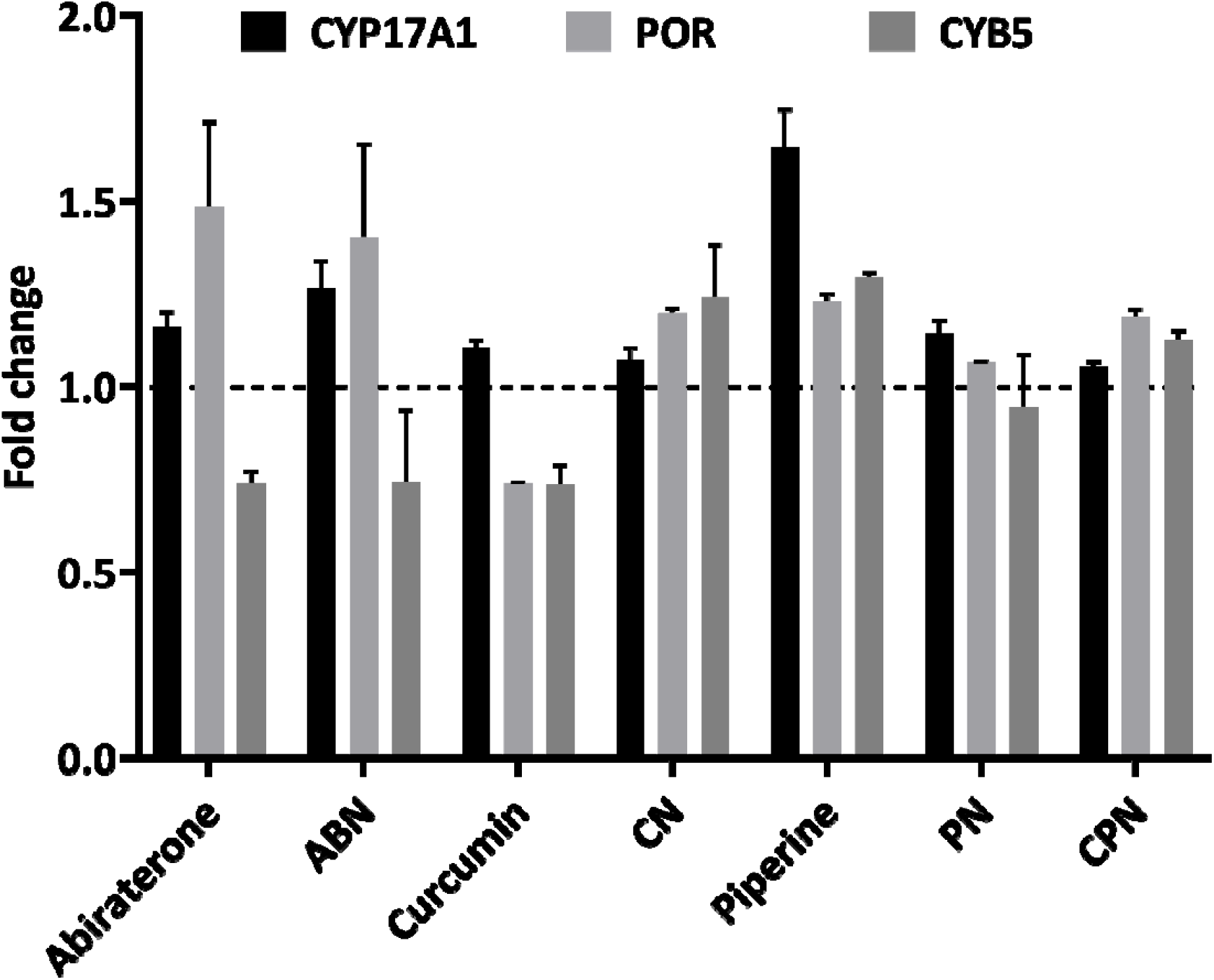
Gene expression profiling for NCI-H295R cells treated with polymer, DMSO, ethanol, abiraterone, ABN (abiraterone nanoparticles), curcumin, piperine, PN (piperine nanoparticles), CN (curcumin nanoparticles), and CPN (curcumin piperine nanoparticles) by qRT-PCR. Total RNA was isolated from NCI-H295R cells grown in normal growth medium. The GAPDH gene was used as internal control for data normalization. Analysis of relative gene expression values was performed by the 2−ΔΔCt method. Summaries of qRT-PCR results of three independent experiments (mean⍰±⍰SD) are shown. * p⍰< ⍰0.05, ** p⍰< ⍰0.01.

### POR Expression

POR exhibited varied responses to the treatments. Abiraterone resulted in an increase in POR expression, with values ranging from 1.32 to 1.64-fold of control (Figure 8). This increase may reflect a compensatory mechanism to support enhanced electron transfer due to CYP17A1 inhibition. The nanoparticle formulations of curcumin and piperine showed minimal impact on POR expression, with values ranging from 1.19 to 1.17-fold and 1.06 to 1.07-fold of control, respectively. This suggests that these treatments do not significantly alter POR expression. Conversely, curcumin alone decreased POR expression to 0.74-fold of control, while piperine alone had a minimal effect, changing POR expression to 1.24 and 1.22-fold of control.

### CYB5A Expression

Abiraterone caused a modest decrease in CYB5 expression, ranging from 0.71 to 0.76-fold of control (Figure 8). This could be a compensatory response to the inhibition of CYP17A1. The nanoparticle formulations of curcumin and piperine changed CYB5 expression to 1.14 to 1.34-fold and 1.04 to 1.29-fold of control, respectively, indicating a significant impact on steroidogenic pathways through reduced availability of electron transfer proteins. Curcumin alone showed a slight decrease in CYB5 expression to 0.77 and 0.70-fold of control, while piperine alone showed a moderate increase to 1.30 and 1.29-fold of control. These effects align with the observed changes in CYP17A1 expression.

Our results shows that curcumin and piperine, especially in nanoparticle formulations, modulate the expression of key steroidogenic enzymes, including CYP17A1. These results prove their potential therapeutic benefits in targeting steroid biosynthesis pathways, which could be relevant for PC where androgen levels are critical.

### Effects of Various Treatments on Cell Cycle Regulation in LNCaP Cells

We investigated the effects of several compounds, including DMSO, ethanol, polymer, abiraterone, curcumin, CN (curcumin nanoparticles), piperine, PN (piperine nanoparticles), and CPN (curcumin piperine nanoparticles), on the cell cycle phases (G2/M, G0/G1, and S phase) in LNCaP cells using FACS analysis. The cells were treated with these compounds for 24 hours, and their impact on cell cycle distribution was analyzed (Figure 9). The gated pictures for the all the treatment is represented as Supplementary Figure S2.

**Figure 9:**
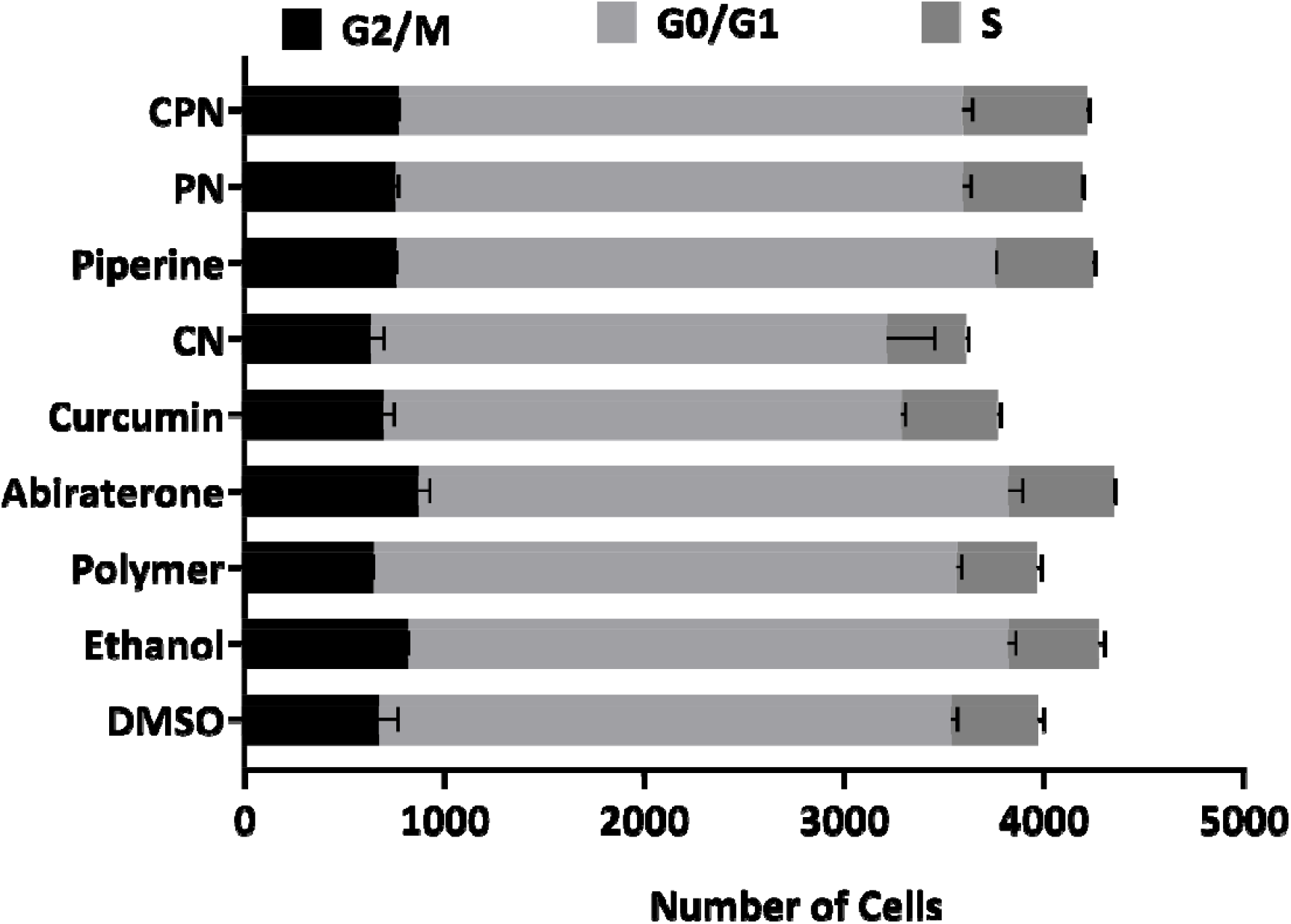
LNCaP cells were treated with DMSO, ethanol, polymer, abiraterone, curcumin, curcumin nanoparticles (CN), piperine, piperine nanoparticles (PN), and curcumin piperine nanoparticles (CPN) for 24 hours. After treatment, cells were harvested, and DNA content was analyzed by propidium iodide staining followed by FACS to assess cell cycle distribution. Quantitative analysis represents the mean percentage distribution of cells in G0/G1, S, and G2/M phases from three independent experiments. Error bars indicate mean ± SD.

### G2/M Phase

Abiraterone treatment resulted in a notable increase in the G2/M phase cell population (913 and 832), suggesting that abiraterone may induce cell cycle arrest in the G2/M phase. This is consistent with the known mechanisms of abiraterone, which targets androgen biosynthesis pathways critical for cell proliferation in PC. Piperine and PN also increased the G2/M phase cell population (763 and 759 for Piperine; 768 and 741 for PN), indicating potential cell cycle arrest effects.

### G0/G1 Phase

Curcumin and its nanoparticle formulation (CN) led to a decrease in the G0/G1 phase cell population (2576 and 2606 for Curcumin; 2751 and 2412 for CN), indicating potential cell cycle arrest effects. Curcumin’s impact on the cell cycle could be linked to its ability to inhibit key steroidogenic enzymes such as CYP17A1, which is crucial for androgen biosynthesis. Abiraterone, despite increasing the G2/M phase, did not significantly alter the G0/G1 phase (2895 and 3003), suggesting a specific G2/M arrest.

### S Phase

The combination treatment with CPN showed the highest increase in the S phase cell population (631 and 614), suggesting enhanced S phase entry or prolonged S phase duration. This effect could be attributed to the synergistic interaction between curcumin and piperine, enhancing their combined efficacy in disrupting cell cycle progression. The increased S phase population might indicate enhanced DNA replication stress or prolonged DNA synthesis phases, which is a hallmark of effective anticancer strategies. Abiraterone also increased the S phase population (526 and 533), supporting its role in cell cycle disruption.

Our results suggest that the various treatments differentially affect cell cycle progression in LNCaP cells. Abiraterone’s induction of G2/M arrest and increased S phase population aligns with its role in inhibiting androgen biosynthesis, thereby affecting cell proliferation. Curcumin and its nanoparticle formulations, along with piperine, demonstrate significant effects on cell cycle regulation, potentially through their inhibitory actions on key steroidogenic enzymes. These findings align with previous studies that have demonstrated curcumin’s ability to interfere with cell cycle regulators, such as cyclins and cyclin-dependent kinases (CDKs), and its potential to induce cell cycle arrest in cancer cells^74^. The enhanced effect observed with nanoparticle formulations, particularly the combination of curcumin and piperine, underscores the potential of these formulations to improve the bioavailability and efficacy of curcumin in cancer therapy.

Our investigation demonstrates that curcumin and piperine, particularly when delivered through nanoparticle formulations, exhibit substantial promise as therapeutic agents in PC. The enhanced cellular uptake and bioavailability of CPN, as evidenced by fluorescence microscopy, translate into superior anticancer effects, as observed in the cell viability and wound healing assays. The ability of CPN to significantly inhibit key steroidogenic enzymes like CYP17A1 suggests a targeted disruption of androgen biosynthesis, critical for PC progression. Furthermore, the modulation of cell cycle phases, particularly the increased S phase population, underscores the potential of these formulations to impact cancer cell proliferation. The selective toxicity towards cancer cells and the sparing of normal cells highlights the specificity and reduced off-target effects of CPN. Overall, the combined effects of curcumin and piperine in nanoparticle form provide a promising approach for enhancing PC treatment and potentially improving patient outcomes. Future studies should explore the clinical translation of these findings and assess long-term efficacy and safety in broader therapeutic contexts.

## Materials and methods

### Materials

Trilostane was extracted from commercially available Modrenal® tablets (Bioenvision, New York, USA). Abiraterone acetate was procured from MedChemExpress^®^ through Lucerna Chem AG (Lucerne, Switzerland). Radiolabeled substrates, including progesterone [4-14C] (specific activity: 55 mCi/mmol; concentration: 0.1 mCi/mL), 17α-hydroxypregnenolone [21-3H] (specific activity: 15 Ci/mmol; concentration: 1 mCi/mL), and [3H]-17α-hydroxy progesterone [1, 2, 6, 7-3H] (specific activity; 60-120 Ci/mmol; 2.22-4.44 TBq/mmol), were sourced from American Radiolabeled Chemicals Inc. (St. Louis, MO, USA).

Non-radiolabeled substrates such as pregnenolone, progesterone, 17α-hydroxyprogesterone, 17α-hydroxy pregnenolone, Resazurin sodium salt, and dimethyl sulfoxide (DMSO) were obtained from Sigma-Aldrich^®^ (St. Louis, MO, USA). Organic solvents were acquired from Carl Roth^®^ GmbH + Co. KG (Karlsruhe, Germany), while activated charcoal was sourced from Merck AG (Darmstadt, Germany). Silica gel-coated aluminum backed thin-layer chromatography (TLC) plates were purchased from Macherey-Nagel (Oensingen, Switzerland). The tritium screens used for autoradiography were obtained from Fujifilm (Dielsdorf, Switzerland). Turmeric extract capsules (Curcuma longa) were sourced from the Finest Natural (Item: 943.17, Deerfield, IL, USA).

Resomer^®^ RG 502 H, poly (D, L-lactide-*co*-glycolide) with lactide:glycolide ratio of 50:50 (Mw 7,000-17,000 g/mol), poly (vinyl alcohol) (PVA) (Mw 9,000-10,000 g/mol, 80% hydrolyzed), and acetonitrile were obtained from Sigma-Aldrich (St. Louis, MO, USA).

### Cell Lines and Culture Media

PC cell lines LNCaP (FGC CRL-1740) were purchased from the American Type Culture Collection (ATCC^®^) and cultured in Roswell Park Memorial Institute (RPMI)-1640 Medium containing 2 mM L-glutamine, 10 mM HEPES, 1 mM sodium pyruvate, 10% fetal bovine serum (FBS), and 1% penicillin-streptomycin (Gibco^™^, Thermo Fisher Scientific, Waltham, MA, USA). VCaP (CRL-2876), PC-3 (CRL-1435), DU-145 (HTB-81), and RWPE-1 (CRL-3607) were provided by Prof. Mark Rubin from the Department of Biomedical Research (DBMR) at the University of Bern, Switzerland. PC-3 was cultured in the same medium as LNCaP cells. VCaP and DU-145 cells were cultured in Dulbecco’s Modified Eagle Medium (DMEM) medium supplemented with 10% FBS, 1% antibiotic mix (100×), and 1 mM sodium pyruvate. RWPE-1 cells were cultured in keratinocyte serum-free medium supplemented with 0.05 mg/mL bovine pituitary extract, 5 ng/mL human recombinant epidermal growth factor, 1% penicillin (100 U/mL), and streptomycin (100 μg/mL; GIBCO). Human adrenocortical NCI-H295R cells (from ATCC^®^: CRL-2128) were grown in DMEM/Ham’s F-12 medium containing L-glutamine and 15 mM HEPES (Thermo Fisher Scientific, Waltham, MA, USA), supplemented with 5% NU-I serum (Becton Dickinson, Franklin Lakes, NJ, USA), 0.1% insulin, transferrin, selenium (100 U/mL; Thermo Fisher Scientific, Waltham, MA, USA), 1% penicillin (100 U/mL; Thermo Fisher Scientific, Waltham, MA, USA), and streptomycin (100 μg/mL; GIBCO). Caco-2 HTB-37 were provided by Dr. Kandasamy Palanivel and were cultured in DMEM supplemented with 0.1 mM of nonessential amino acids, 100 U/mL of penicillin, 0.1 g/mL of streptomycin, 10 mM of sodium bicarbonate, and 10% FBS. Passage numbers during the experiments remained below 30 according to established protocols^8,9,13,64,75^.

### Curcuminoid Extraction and Preparation of Nanoparticles

The curcuminoids were extracted, purified, and quantified following the methodology previously described [38]. PLGA nanoparticles were prepared using the nanoprecipitation technique (Figure 10) [44, 63-65]. The organic phase containing a mixture of 10 mg PLGA and 2 mg Curcumin or Piperine was prepared in 500 μL acetonitrile. For the PLGA nanoparticles co-loaded with Curcumin and Piperine, 1 mg of each drug was used to keep the same drug: polymer ratio. The organic phase was added dropwise to the 2 mL of aqueous phase containing 2% PVA under rigorous stirring. The solution was kept stirring overnight at room temperature, in the dark, to evaporate the organic solvent. The nanoparticles were then washed by three centrifugation cycles (15000 rpm, 35 min.), the supernatant was discarded, and the pellet was resuspended in deionized water for characterization and usage. The nanoparticles generated included Piperine nanoparticle (PN), Curcumin nanoparticle (CN), Curcumin-Piperine nanoparticle (CPN) and the control nanoparticle (contains no drug).

**Figure 10:**
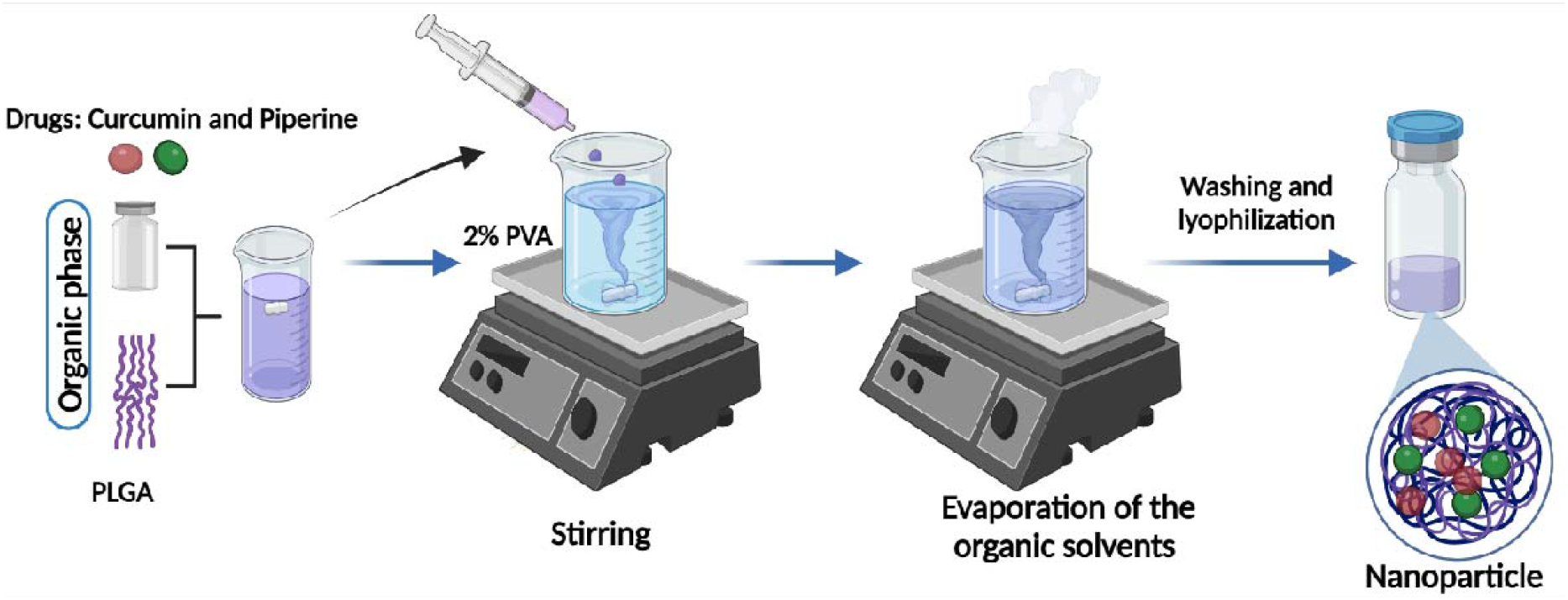
Scheme for the nanoparticle preparation. The organic phase containing PLGA, and drugs was added dropwise to the aqueous phase composed of 2% PVA. Nanoparticles were washed and lyophilized following the evaporation of the organic solvent. Created with biorender.com.

### Characterization of the Nanoparticles

The hydrodynamic size and polydispersity index (PDI) of nanoparticles were measured by dynamic light scattering (DLS) using a Zeta Sizer Nano ZS (Malvern, UK) in the backscattering mode. 1 mg/mL of particle suspensions were prepared in water and vortexed prior to measurements. Each sample underwent size analysis three times, with 10 runs each. Hydrodynamic size and PDI of the nanoparticles were reported as average ± standard deviation.

The morphology and surface topography of the nanoparticles were characterized using a multi-mode NanoScope atomic force microscope (AFM) (Bruker). 100 μL of 1 mg/mL particle suspension was dried on clean glass substrates and particles were imaged in peak-force tapping mode. Silicon nitride cantilevers with nominal spring constants of 0.4 N/m (Bruker) were used for the imaging. The scan rate was set at 1 Hz. 256 lines with 256 points per line were recorded during image acquisition. AFM images were analyzed using NanoScope analysis software (Bruker).

To evaluate the drug encapsulation efficiency, a previously described method ^35^ with a few modifications was employed. Two milligrams of nanoparticles were dissolved in 1 mL of methanol and gently shaken at 4 °C for 36 hours to ensure complete leaching of curcumin and piperine from the nanoparticles into the methanol. The resulting solutions were centrifuged at 14000 rpm, and the supernatants were collected for analysis. The concentrations of curcumin and piperine in the 1:50 dilutions were determined using a UV-Vis spectrophotometer, with absorbance measurements taken at 342 nm for piperine and 450 nm for curcumin. Standard calibration curves for piperine and curcumin (ranging from 0 to 20 μg/mL) were prepared under identical conditions to quantify the drug concentrations.

### Fluorescence Microscopy

Sterile coverslips were placed into 6-well plates and seeded with Caco-2 cells at a density of 5,000 cells per well in complete medium. The cells were treated with curcumin, CN, and CNP for 4 hours. Previous reports indicated that curcumin exhibits cytosolic staining, which can be attributed to encapsulated curcumin rather than its free form ^76-78^. Following treatment, the coverslips were washed in sodium phosphate buffer, air-dried, and mounted using ProLong^®^ Gold Antifade Reagent with DAPI (Invitrogen, Grand Island, NY, USA), then stored at 4°C. The cells were visualized using Leica DM5500B widefield microscope equipped with an LCD camera at 20× magnification. Excitation and emission wavelengths of 420 nm and 540 nm, respectively, were used to observe the stained cells.

### Measurement of Curcumin Uptake

CN and CPN were administered to NCI-H295R cells (5 × 10^5^ cells/well) cultured in 6-well plates and incubated at 37°C with 5% CO_2_ and 90% humidity for 48 hours. Subsequently, the cells were exposed to a new medium containing 20 μM curcumin, CN, and CPN for 4 hours. After incubation, the drug-containing medium was removed, and the cells were washed and recovered using 0.05% trypsin-EDTA. The cells were then centrifuged for 5 minutes at 1,500 rpm at 4°C. The supernatant was discarded, and the pellet was allowed to dry at room temperature in the dark. Once fully dried, the pellet was resuspended in 0.25 mL of methanol and sonicated at 30% amplitude for 5 minutes to extract curcumin from the cells. The mixture was then centrifuged for 5 minutes at 10,000 rpm at 4°C, and the supernatant was collected. The sample was diluted 1:1 in methanol and analyzed using a UV/Visible spectrophotometer as described by ^13,79^.

### Cell Viability Assay Using Resazurin Sodium Salt

NCI-H295R, LNCaP, PC-3, DU-145, and RWPE-1 cells were seeded into 96-well culture plates at a density of 1 × 10^4^ cells per well and incubated overnight at 37°C with 5% CO_2_ and 90% humidity. VCaP cells were seeded at a density of 1.2 × 10^4^ cells per well under the same conditions. After 24 hours, the medium was replaced, and PN, CN, CPN, Piperine, curcumin, and abiraterone the drugs were added to the medium at a final concentration of 10 µM. The cells were then incubated for an additional 24 and 48 hours. Cell viability was assessed using the Alamar Blue assay ^49,75^. Post-incubation, 0.05 mg/mL Alamar Blue in phosphate-buffered saline (PBS) was added to each well. The plates were incubated for 4 hours in the dark at 37°C, and fluorescence was measured at an excitation wavelength of 550 nm and an emission wavelength of 590 nm. Percent viability was calculated relative to the mean value of the control samples treated with DMSO, NN and Ethanol. Additionally, the cytotoxicity of curcuminoids on human HEK-293 cells at a density of 1 × 10^4^ cells per well was evaluated under similar conditions using the same assay. All experiments were performed in triplicate.

### Wound-Healing Assay

DU-145 cells were seeded in 24-well plates at a density of 50,000 cells per well and cultured until they reached over 85% confluence. The cells were then treated with 5 µg/mL mitomycin for 2 hours to inhibit cell proliferation. A wound was created using a 10 µL pipette tip. Subsequently, the cells were treated with drugs at a concentration of 10 µM, while control wells were treated with DMSO, or ethanol. Scratch closure (Wound healing) was monitored and imaged using an optical microscope after 24 h. The resulting images were analyzed with ImageJ software (version 1.54j) ^80^.

### Assay of CYP17A1 and CYP21A2 Activities

NCI-H295R cells were seeded in 12-well plates at a density of 500,000 cells per well and incubated overnight. The following day, 10 µM of test compounds were added to the wells with fresh medium and incubated for 4 hours. Abiraterone served as a reference, while DMSO, NN, and ethanol were used as controls. To assess CYP17A1 hydroxylase activity, cells were treated with [^14^C]-Progesterone at a concentration of 10,000 cpm/1 µM per well. Trilostane was added prior to the test compounds and the substrate to inhibit 3β-hydroxysteroid dehydrogenase activity. Radiolabeled steroids were extracted from the media using an ethyl acetate and isooctane mixture (1:1 v/v) and separated via TLC on silica gel-coated aluminum plates (Supelco^®^ Analytics, Sigma-Aldrich Chemie GmbH, Taufkirchen, Germany). TLC spots were exposed to a phosphor screen and detected by autoradiography using a Typhoon^™^ FLA-7000 PhosphorImager (GE Healthcare, Uppsala, Sweden). Radioactivity was quantified with ImageQuant^™^ TL analysis software (GE Healthcare Europe GmbH, Freiburg, Germany), and enzyme activity was calculated as the percentage of radioactivity incorporated into the product relative to the total radioactivity.

For CYP17A1 lyase activity, cells were treated with 50,000 cpm/1 µM [21-3H]-17α-hydroxypregnenolone per well under similar conditions. The tritiated water release assay measured the conversion of 17α-hydroxypregnenolone into DHEA. Steroids in the media were precipitated using a 5% activated charcoal/0.5% dextran solution. Enzyme activity was estimated based on the water-soluble tritiated by-product formed in an equimolar ratio with DHEA, with radioactivity in the aqueous phase measured by liquid scintillation counting (MicroBeta2^®^ Plate Counter, PerkinElmer Inc., Waltham, MA, USA). Percent inhibition was calculated relative to the control.

To determine cytochrome P450 21-hydroxylase (CYP21A2) activity, [^3^H]-17α-hydroxyprogesterone (∼50,000 cpm/1 µM per well) was used as a substrate. Following incubation, the medium from each well was collected, and steroids were extracted as previously described for CYP17A1 hydroxylase activity ^9,13,64,75^.

### Steroid Profiling

Steroid levels were quantified using liquid chromatography–high-resolution mass spectrometry (LC-HRMS), following previous studies ^75,81^. NCI-H295R cells were seeded in 12-well plate at a density of 500,000 cells per well and treated with fresh growth media containing the test drugs for 24 hours. Subsequently, 1 µM of pregnenolone was added and incubated for an additional 4 hours. Steroids were then extracted from 500 µL aliquots of cell media, separated, and quantified according to the protocol described by ^81^.

### Quantitative Real Time PCR (qRT-PCR)

NCI-H295R cells were treated with compounds for 24 hours, after which total RNA was extracted using the TRIzol reagent following the manufacturer’s protocol (Invitrogen, Carlsbad, CA, USA). RNA was then reverse transcribed into complementary DNA (cDNA) utilizing the Improm II Reverse Transcriptase kit (Promega, Madison, WI, USA). Quantitative reverse transcription PCR (qRT-PCR) was performed on the 7500 Fast Real-Time PCR System (Applied Biosystems, Foster City, CA, USA) using ABsolute SYBR Green Mix (ABgene, Thermo Fisher Scientific, Waltham, MA, USA). GAPDH served as the endogenous control for normalization. Gene expression levels were quantified using the 2^−ΔΔCt^ method to determine fold changes. Amplification curves and mean cycle threshold (Ct) values were analysed with the 7500 Fast System Software (Applied Biosystems). The ΔCt and ΔΔCt values were calculated as previously described ^82,83^.

### Cell Cycle Analysis

For cell cycle analysis, LNCaP cells were cultured under normal growth conditions and treated with compounds. After a 24-hour incubation period, cells were harvested and resuspended in 400 μL of a solution containing 50 μg/mL propidium iodide (PI) in a buffered medium. Propidium iodide intercalates with DNA and allows for the measurement of DNA content. Stained cells were analysed using the Amnis ImageStreamX Mark II flow cytometer (Amnis Corporation, Seattle, WA, USA), which provides high-resolution images of individual cells and enables detailed cell cycle profiling. Quantitative analysis of cell cycle phases was performed using the IDEAS software (Amnis Corporation, Seattle, WA, USA), which facilitates precise measurement of fluorescence intensity and cell cycle distribution based on the DNA content ^83^.

### Statistical Analysis

For statistical analysis, R Studio (version 3.6.0+) and GraphPad Prism v8.0 (GraphPad Software, Inc., San Diego, CA, USA) were utilized to enable data evaluation. Data are presented as the mean ± standard deviation (SD) of three independent replicates to account for variability and reproducibility. One-way analysis of variance (ANOVA) was employed to assess differences between the treatment groups and their corresponding controls, providing a robust statistical framework for detecting significant variations. Post-hoc analyses were performed using Tukey’s Honest Significant Difference (HSD) test to identify specific group differences. Significance thresholds were set at *p < 0.05 and **p < 0.001, All statistical tests were two-tailed, and assumptions of normality and homogeneity of variances were verified prior to analysis to ensure the validity of the ANOVA results.

## Conflict of Interest statement

The authors declare no conflict of interest.

## Acknowledgement

We thank Prof. Mark Rubin for providing the RWPE1, DU145, VCaP, and PC3 cell lines and Dr. Kandasamy Palanivel for the Caco2 cells. The authors are grateful to Kojo Atchou for help with the fluorescence microscopy.

## Author Contributions

Conceptualization, A.V.P., CYP17A1 assays, J.Y.; cell viability, I.S.B., J.Y.; steroid profiling, T.d.T.; preparation and characterization of nanoformulations, E.N., J.Y., O.T.; writing— original draft preparation, JY.; writing—review and editing, JY, T.d.T., A.V.P., O.T.; project administration AVP. All authors have read and agreed to the published version of the manuscript.

## Data Availability Statement

Data are available in manuscript text or in supplementary materials.

## Funding statement

This research was supported by CANCER RESEARCH SWITZERLAND grant number KFS-5557-02-2022 to A.V.P. J.Y., is. partially funded by the SWISS GOVERNMENT EXCELLENCE SCHOLARSHIP (ESKAS) grant number 2022.0470, and. T.d.T. was funded by a Marie Skłodowska-Curie Individual Fellowship (#101023999).

## Notes

### Competing Interest Statement

The authors have declared no competing interest.

