## Supplementary Data for "Nanoparticles with Curcumin and Piperine Modulate Steroid Biosynthesis in Prostate Cancer"

Supplementary Table S1. The cell viability results of the drugs on non-prostate cancer cell lines.

| Cell viability results of the drugs on HEK293T cells | | | | | | | | | | |
|---|---|---|---|---|---|---|---|---|---|---|
| | **24 hours** | | | | | **48 hours** | | | | |
| **Drugs** | **%Control** | | | **Mean** | **SD** | **%Control** | | | **Mean** | **SD** |
| **DMSO** | 99.9153 | 100.501 | 99.5841 | 100 | 0.46413 | 93.3076 | 98.3343 | 108.358 | 100 | 7.66223 |
| **Polymer [Empty nanoparticle]** | 100.332 | 100.107 | 99.561 | 100 | 0.39637 | 92.4785 | 105.185 | 102.336 | 100 | 6.66775 |
| **Ethanol** | 101.265 | 98.337 | 100.398 | 100 | 1.50397 | 109.019 | 88.603 | 102.378 | 100 | 10.4136 |
| **Abiraterone** | 133.215 | 130.787 | 130.716 | 131.573 | 1.42229 | 121.044 | 129.033 | 121.796 | 123.958 | 4.41148 |
| **Abiraterone nanoparticle** | 111.307 | 111.743 | 112.057 | 111.703 | 0.37671 | 86.0048 | 86.8621 | 85.889 | 86.2519 | 0.53157 |
| **Curcumin** | 74.646 | 73.3742 | 72.8 | 73.6067 | 0.94468 | 33.4154 | 38.3812 | 40.7866 | 37.5277 | 3.75896 |
| **Curcumin nanoparticle** | 87.1068 | 88.5939 | 87.0433 | 87.5813 | 0.87751 | 98.7113 | 86.1354 | 86.3333 | 90.3933 | 7.20427 |
| **Curcumin_piperine nanoparticle** | 83.067 | 83.4948 | 81.5627 | 82.7082 | 1.01478 | 98.7357 | 78.2272 | 69.8108 | 82.2579 | 14.8777 |
| **Piperine** | 84.6663 | 81.4508 | 85.5756 | 83.8976 | 2.16719 | 108.929 | 94.4789 | 121.999 | 108.469 | 13.7656 |
| **Piperine nanoparticle** | 122.264 | 122.985 | 123.227 | 122.825 | 0.50116 | 89.969 | 90.9575 | 105.059 | 95.3285 | 8.44131 |

Supplementary Figure S1: The calibration curve of Piperine

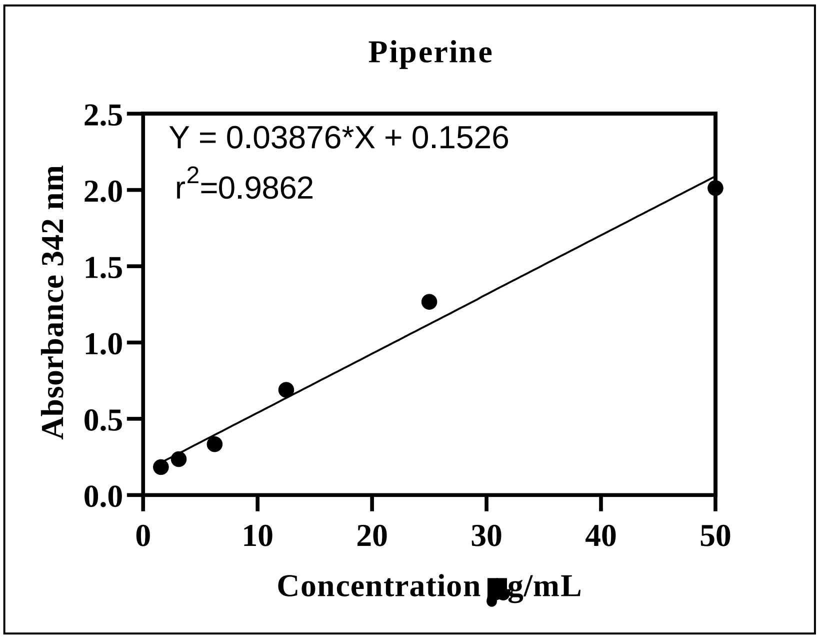

Supplementary Figure S2: Cell cycle gated graphs and sample channel pics

### Abiraterone_treated

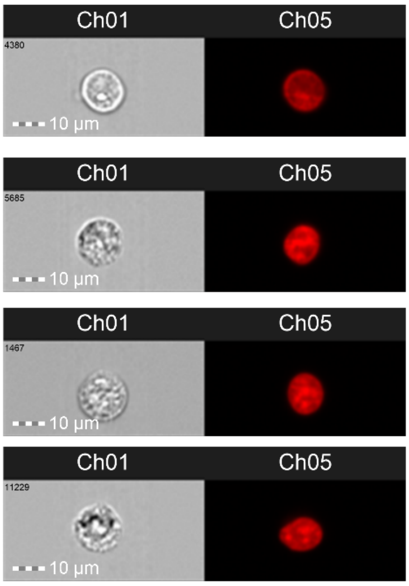


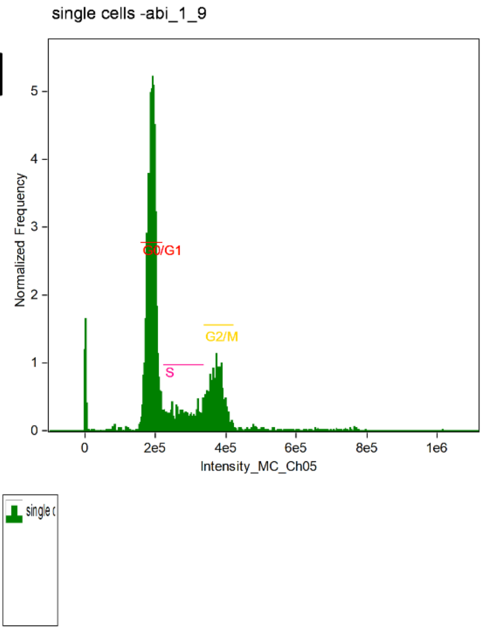


Intensity_MC_Ch05

| Population | Count | %Total | %Gated |
|---|---|---|---|
| single cells & focus | 4676 | 33.5 | 100 |
| G2/M & single cells & focus | 913 | 6.55 | 19.5 |
| G0/G1 & single cells & focus | 2895 | 20.8 | 61.9 |
| S & single cells & focus | 526 | 3.77 | 11.2 |

### Abiraterone nanoparticle_treated

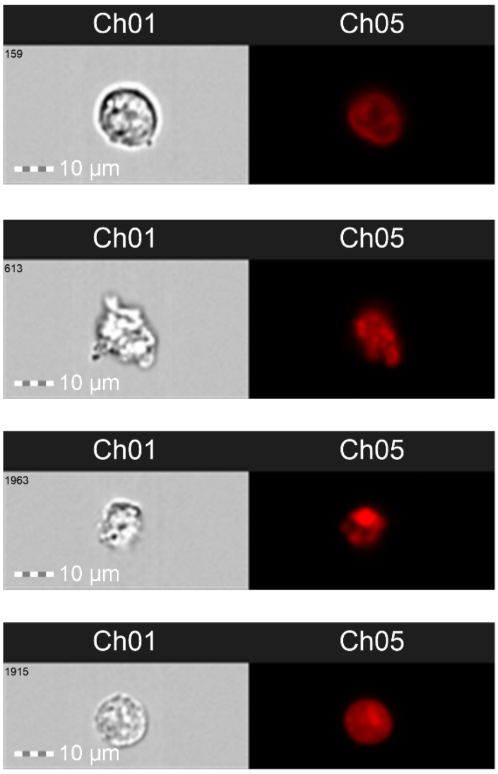


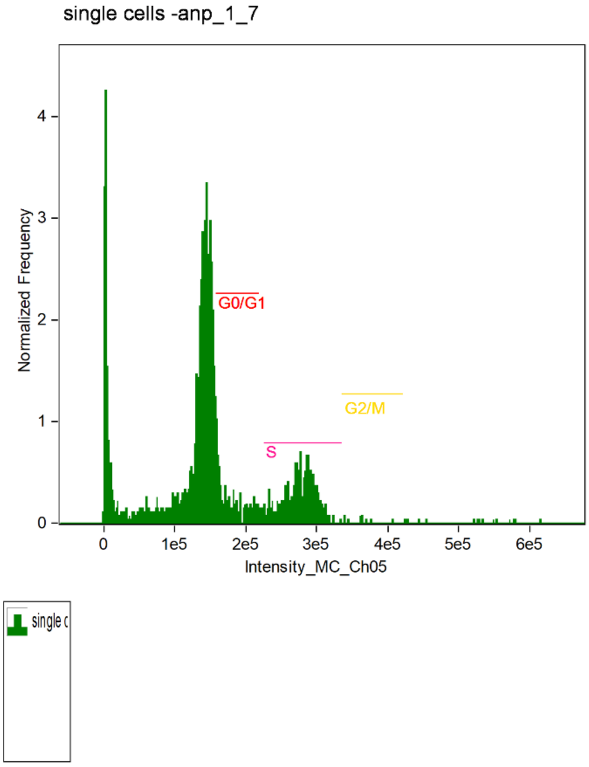


Intensity_MC_Ch05

| Population | Count | %Total | %Gated |
|---|---|---|---|
| single cells & focus | 2723 | 40.2 | 100 |
| G2/M & single cells & focus | 12 | 0.18 | 0.44 |
| G0/G1 & single cells & focus | 259 | 3.83 | 9.51 |
| S & single cells & focus | 473 | 6.99 | 17.4 |

### Curcumin_treated

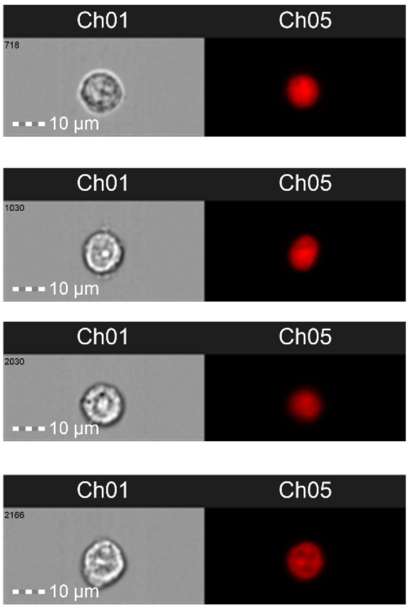


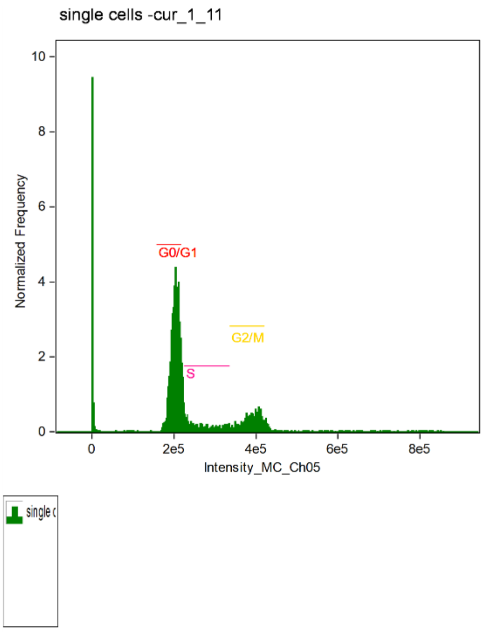


Intensity_MC_Ch05

| Population | Count | %Total | %Gated |
|---|---|---|---|
| single cells & focus | 4891 | 29.8 | 100 |
| G2/M & single cells & focus | 661 | 4.02 | 13.5 |
| G0/G1 & single cells & focus | 2576 | 15.7 | 52.7 |
| S & single cells & focus | 472 | 2.87 | 9.65 |

### Curcumin nanoparticle_treated

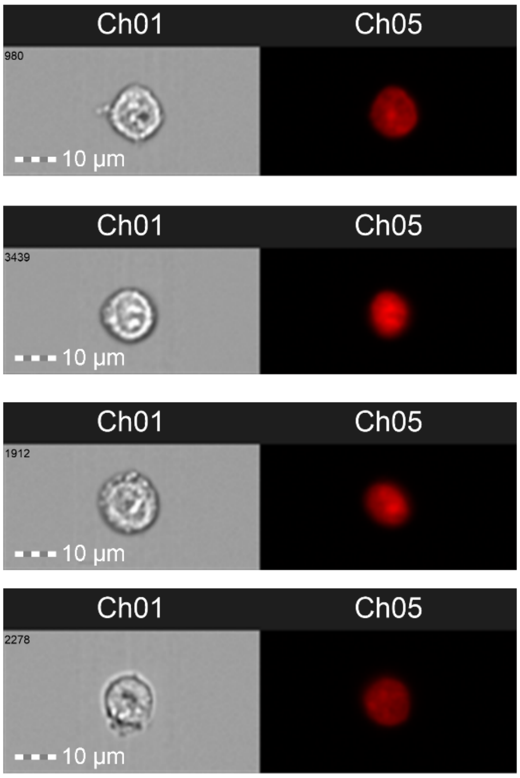


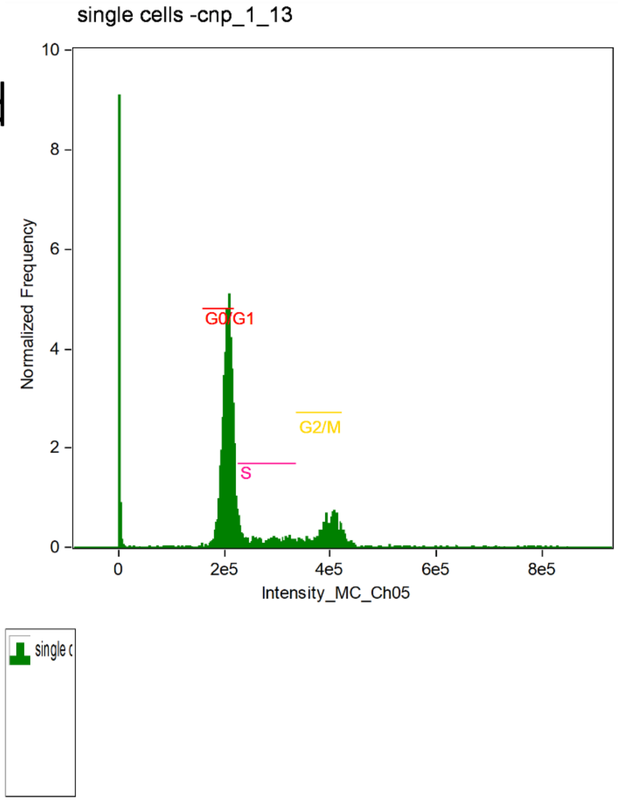


Intensity_MC_Ch05

| Population | Count | %Total | %Gated |
|---|---|---|---|
| single cells & focus | 4526 | 30.8 | 100 |
| G2/M & single cells & focus | 590 | 4.02 | 13 |
| G0/G1 & single cells & focus | 2412 | 16.4 | 53.3 |
| S & single cells & focus | 403 | 2.74 | 8.9 |

### Curcumin_piperine_nanoparticle_treated

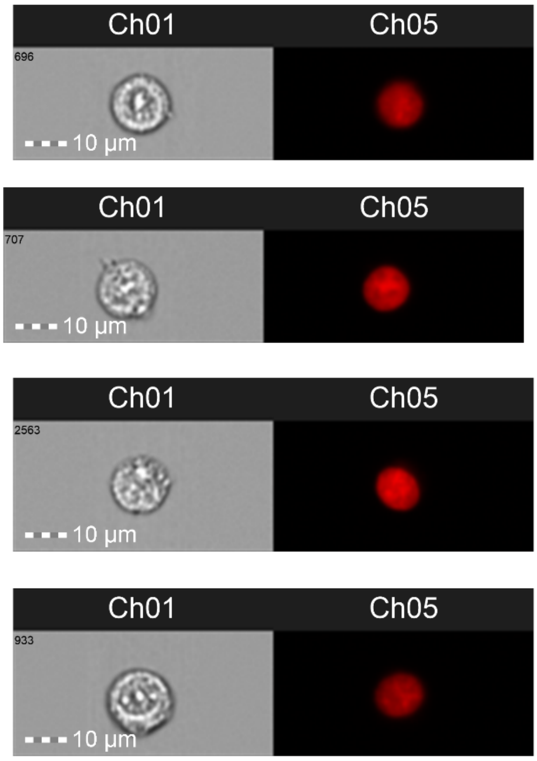


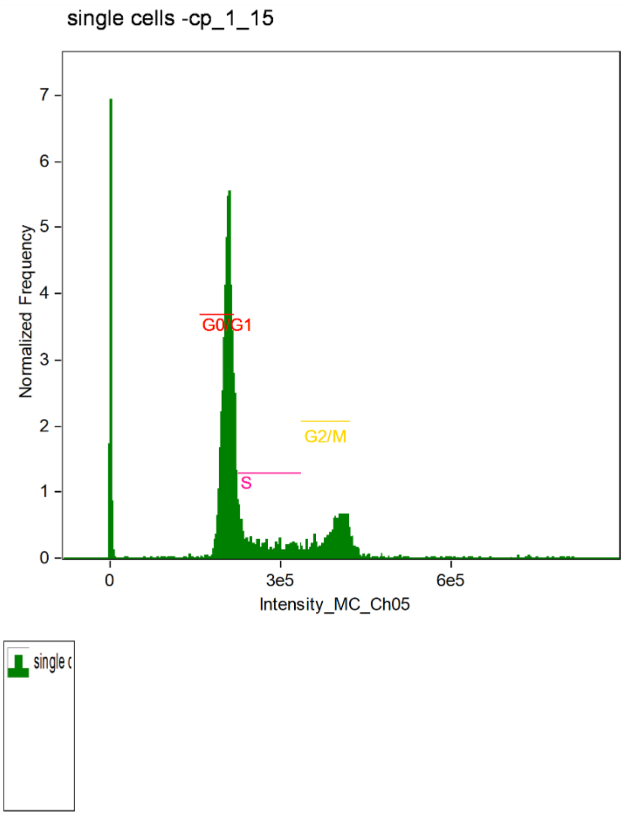


Intensity_MC_Ch05

| Population | Count | %Total | %Gated |
|---|---|---|---|
| single cells & focus | 5098 | 34.5 | 100 |
| G2/M & single cells & focus | 776 | 5.25 | 15.2 |
| G0/G1 & single cells & focus | 2783 | 18.8 | 54.6 |
| S & single cells & focus | 631 | 4.27 | 12.4 |

### Piperine_treated

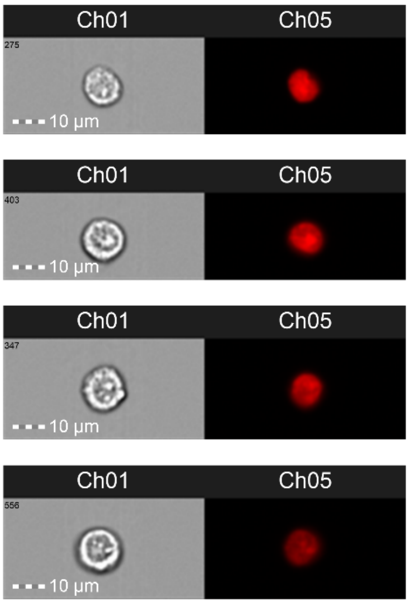


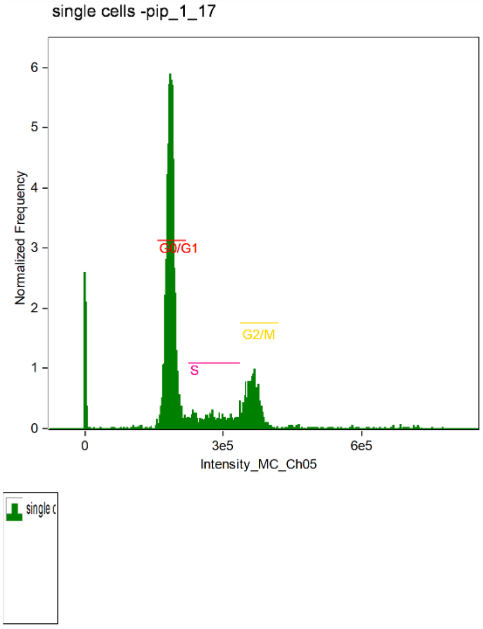


Intensity_MC_Ch05

| Population | Count | %Total | %Gated |
|---|---|---|---|
| single cells & focus | 4625 | 46.4 | 100 |
| G2/M & single cells & focus | 763 | 7.66 | 16.5 |
| G0/G1 & single cells & focus | 3002 | 30.1 | 64.9 |
| S & single cells & focus | 479 | 4.81 | 10.4 |

### Piperine nanoparticle_treated

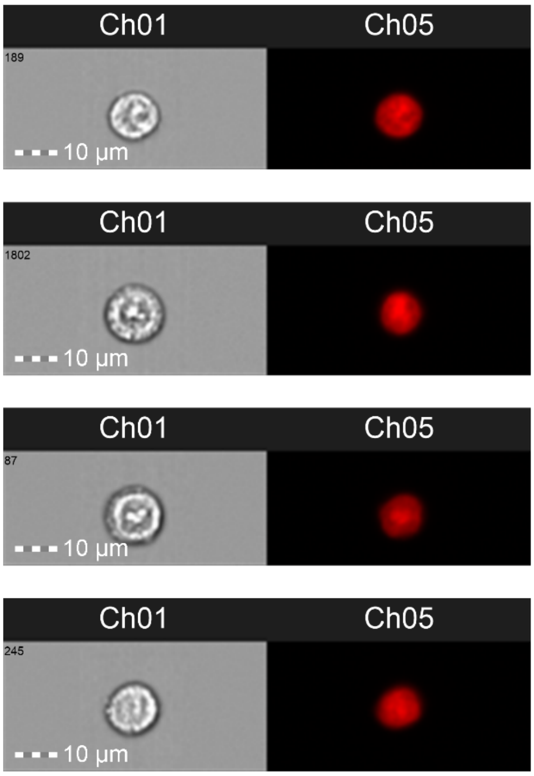


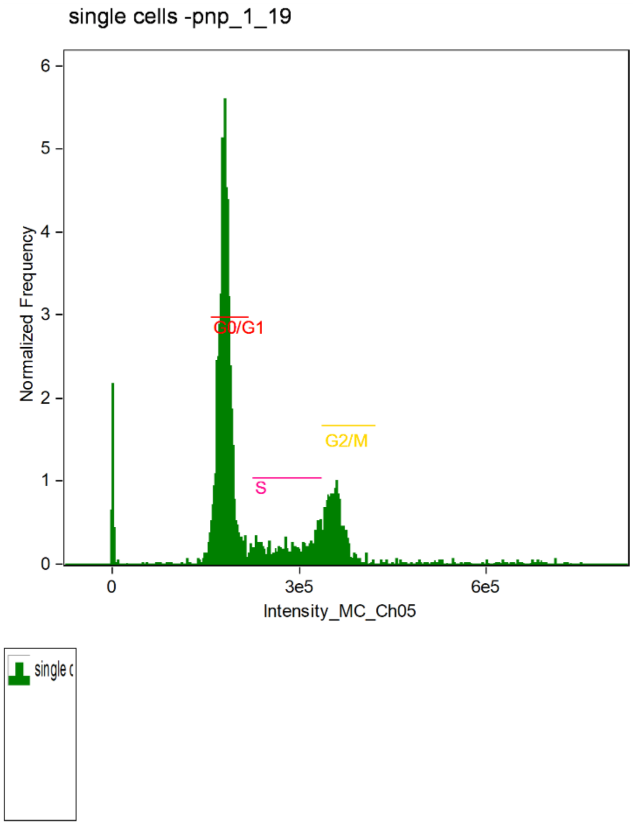


Intensity_MC_Ch05

| Population | Count | %Total | %Gated |
|---|---|---|---|
| single cells & focus | 4594 | 37.4 | 100 |
| G2/M & single cells & focus | 768 | 6.25 | 16.7 |
| G0/G1 & single cells & focus | 2872 | 23.4 | 62.5 |
| S & single cells & focus | 599 | 4.88 | 13 |

### Dmso_treated

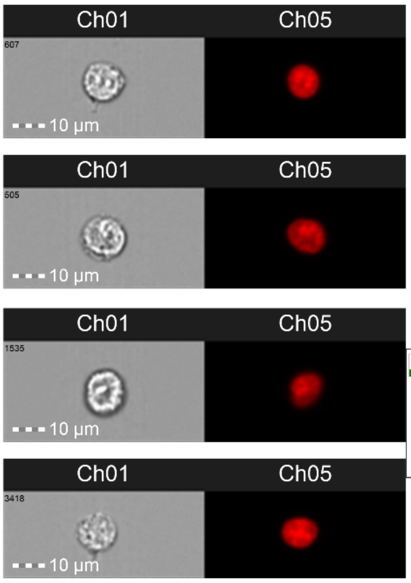


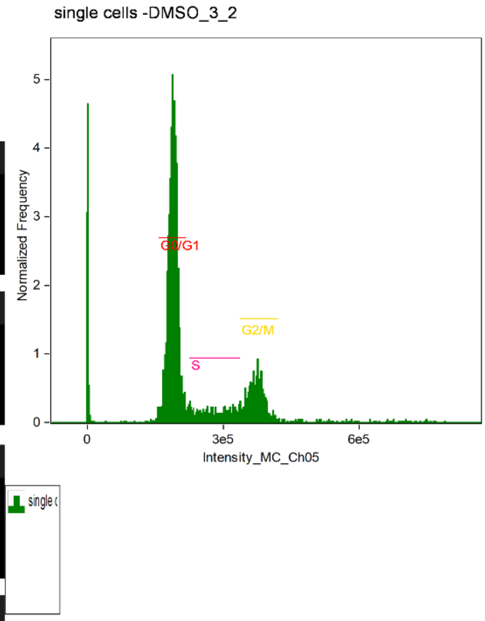


Intensity_MC_Ch05

| Population | Count | %Total | %Gated |
|---|---|---|---|
| single cells & focus | 4550 | 35.8 | 100 |
| G2/M & single cells & focus | 742 | 5.84 | 16.3 |
| G0/G1 & single cells & focus | 2838 | 22.4 | 62.4 |
| S & single cells & focus | 413 | 3.25 | 9.08 |

### Empty nanoparticle [Polymer]_treated

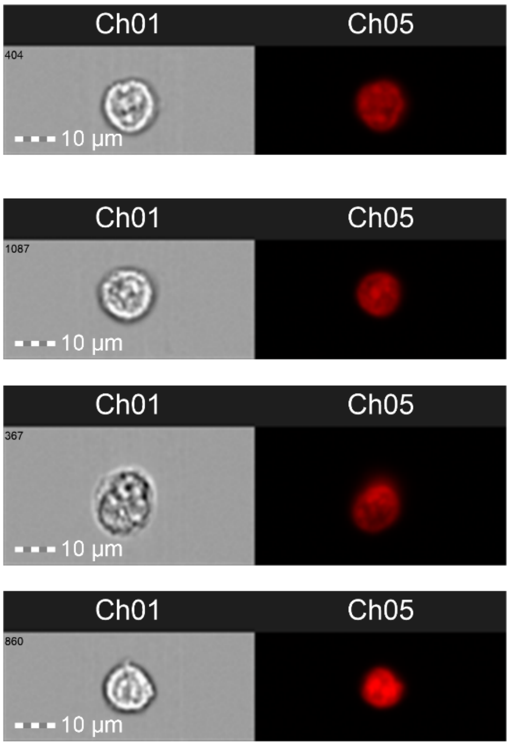


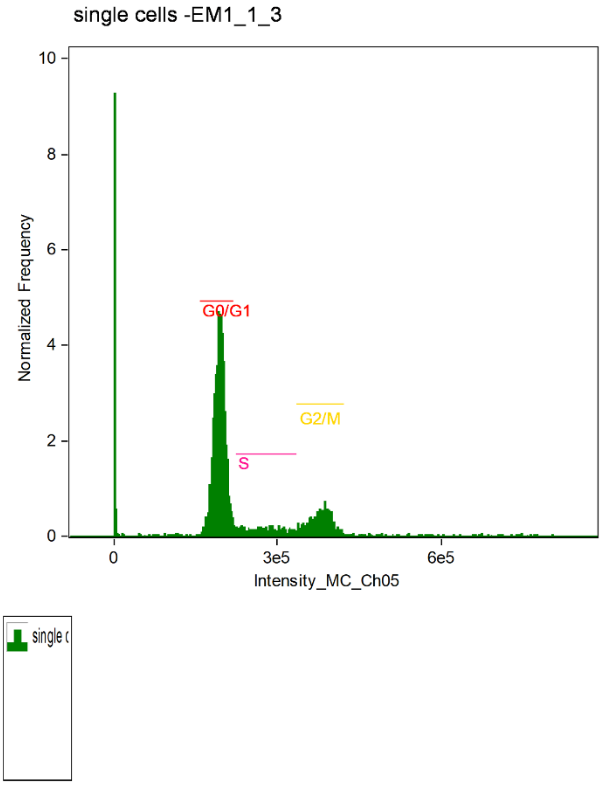


Intensity_MC_Ch05

| Population | Count | %Total | %Gated |
|---|---|---|---|
| single cells & focus | 4852 | 33.7 | 100 |
| G2/M & single cells & focus | 647 | 4.49 | 13.3 |
| G0/G1 & single cells & focus | 2900 | 20.1 | 59.8 |
| S & single cells & focus | 379 | 2.63 | 7.81 |

### Ethanol_treated

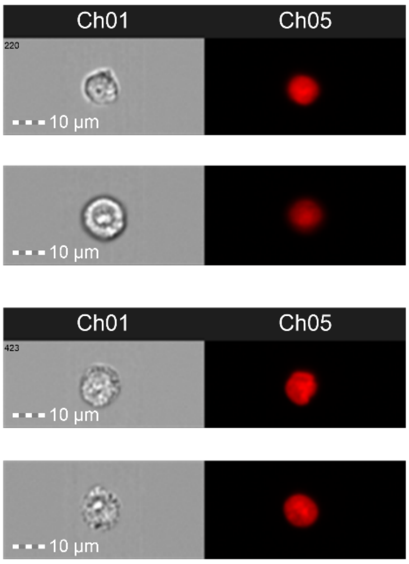


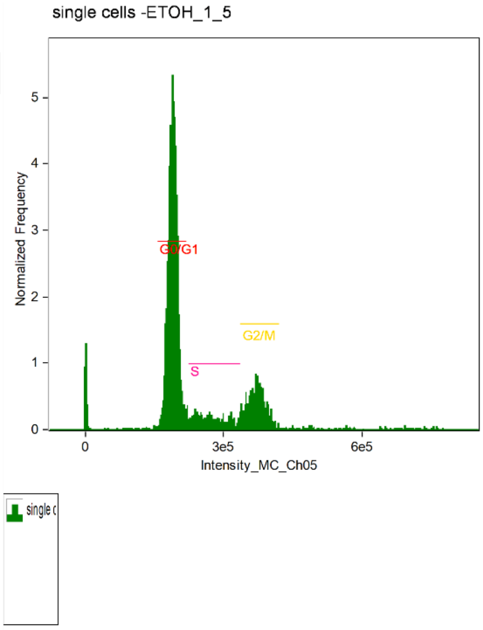


Intensity_MC_Ch05

| Population | Count | %Total | %Gated |
|---|---|---|---|
| single cells & focus | 4565 | 47.2 | 100 |
| G2/M & single cells & focus | 814 | 8.42 | 17.8 |
| G0/G1 & single cells & focus | 3033 | 31.4 | 66.4 |
| S & single cells & focus | 425 | 4.4 | 9.31 |

Supplementary Figure S3: Pictures of scratch assays.

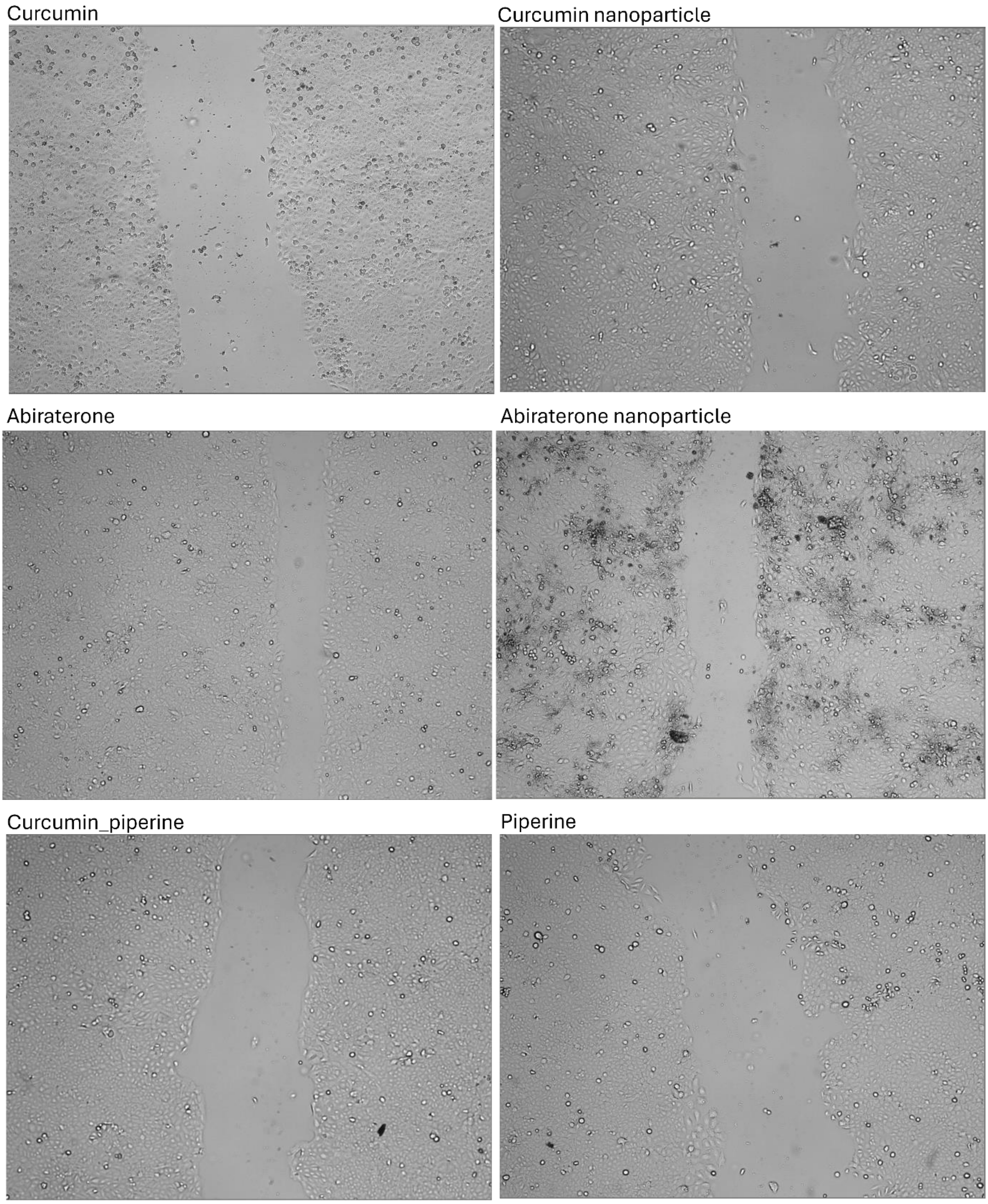

Piperine nanoparticle

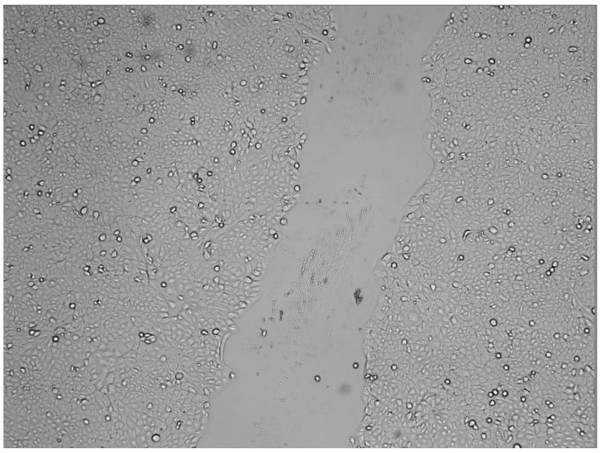

DMSO

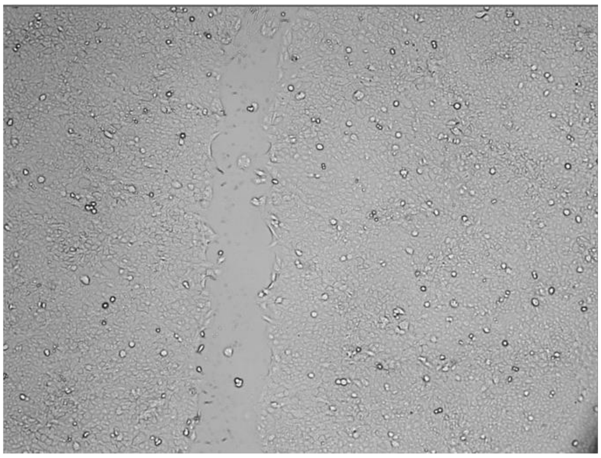

Ethanol

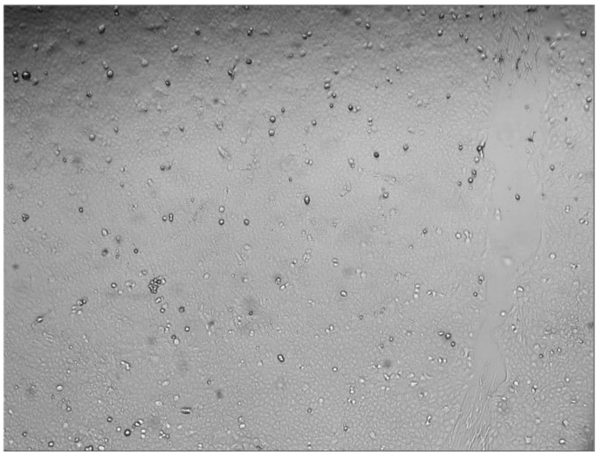

Empty nanoparticle [Polymer]

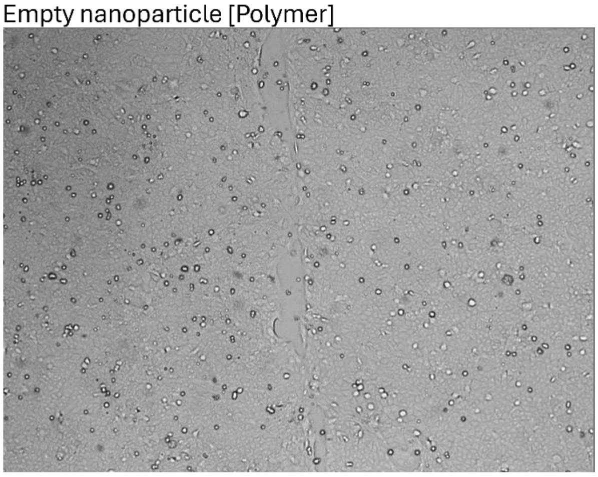
